# Unconventional tonicity-regulated nuclear trafficking of NFAT5 mediated by KPNB1, XPOT and RUVBL2

**DOI:** 10.1101/2021.07.11.451941

**Authors:** Chris Y. Cheung, Ting-Ting Huang, Ning Chow, Shuqi Zhang, Yanxiang Zhao, Catherine C.L. Wong, Daniela Boassa, Sebastien Phan, Mark H Ellisman, John R. Yates, SongXiao Xu, Zicheng Yu, Yajing Zhang, Rui Zhang, Ling Ling Ng, Ben C.B. Ko

## Abstract

NFAT5 is the only known mammalian tonicity-responsive transcription factor functionally implicated in diverse physiological and pathological processes. NFAT5 activity is tightly regulated by extracellular tonicity but the underlying mechanisms remain elusive. We demonstrated that NFAT5 enters the nucleus via the nuclear pore complex. We also found that NFAT5 utilizes a non-canonical nuclear localization signal (NFAT5-NLS) for nuclear imports. siRNA screening revealed that karyopherin β1 (KPNB1) drives nuclear import of NFAT5 via directly interacting with NFAT5-NLS. Proteomics analysis and siRNA screening further revealed that nuclear export of NFAT5 under hypotonicity is mediated by Exportin-T, and that it requires RuvB-Like AAA type ATPase 2 (RUVBL2) as an indispensable chaperone. Our findings have identified KPNB1 and RUVBL2 as key molecules responsible for the unconventional tonicity-regulated nucleocytoplasmic shuttling of NFAT5. These findings offer an opportunity for developing novel NFAT5 targeting strategies that are potentially useful for the treatment of diseases associated with NFAT5 dysregulation.

## Introduction

The nuclear envelope consists of two lipid bilayer membranes that compartmentalize the nucleus from the cytosol. It is perforated with nuclear pore complexes (NPCs) that are composed of proteins known as nucleoporins (Nups). NPCs serve as a gateway to actively control nucleocytoplasmic exchange of proteins and RNAs (D’Angelo and Hetzer, 2008). The control of nuclear availability of transcription factors by nucleocytoplasmic trafficking through NPCs is one of the key mechanisms for regulating gene transcription programs. Under most circumstances, passage of transcription factors across NPCs requires the assistance of specific nucleocytoplasmic transport receptors (NTRs). Proteins in the karyopherin-β (Kapβ) family are the major NTRs in cells that transport most nuclear proteins including transcription factors, and certain classes of RNAs (Kimura and Imamoto, 2014). Kapβs can be further subdivided into the subgroups of nuclear import receptors (importins), export receptors (exportins) and bidirectional receptors depending on their roles in the specific nuclear transport processes. In the human genome, 10 importins, 7 exportins and 2 bidirectional receptors have been discovered. Each of them recognizes a specific protein domain on protein cargos and mediates their passage via NPCs in a Ran GTPase-dependent manner, which provides energy and directionality for the transport process (Pemberton and Paschal, 2005). The best characterized protein domains being recognized by importins and exportins is the classical nuclear localization signals (cNLS) and canonical nuclear export signal (NES), respectively (Lange et al., 2007). Nevertheless, nuclear import signals that do not conform to the cNLS, which are known as nonclassical nuclear localization signals (ncNLS), have also been identified (Soniat and Chook, 2015; Lee et al., 2006b).

Nuclear Factor of Activated T cells (NFAT5), also known as the Osmotic Response Element Binding Protein (OREBP) or Tonicity Responsive Element-binding Protein (TonEBP), is a rel-homology domain containing protein that belongs to the NFAT family, and is the only known tonicity-dependent transcription factor in mammals (Lopez-Rodríguez et al., 1999; Ko et al., 2000; Miyakawa et al., 1999). NFAT5 is indispensable for cell survival under hypertonic milieu, where it restores cellular homeostasis by orchestrating a genetic program that promotes expression of heat shock proteins and replaces detrimental cellular electrolytes by cell compatible organic osmolytes(Burg et al., 2007). Kidney inner medulla was once considered the only physiological relevant site for the activation and activity of NFAT5. Nevertheless, recent findings have suggested the presence of hypertonic stress in certain tissue microenvironments (Go et al., 2004; Machnik et al., 2009), and NFAT5 activation has been associated with a plethora of physiological and pathological processes, including blood pressure regulation, inflammation, and development of autoimmune diseases (Machnik et al., 2009; Kleinewietfeld et al., 2013).

NFAT5 interacts with cognate enhancer, known as Osmotic Response Element (ORE) or Tonicity Responsive Element (TonE) (Takenaka et al., 1994; Ko et al., 1997), for gene transcriptions (Ko et al., 2000; Lopez-Rodríguez et al., 2001; Ferraris et al., 2002). Its activity is controlled by intricate, yet elusive, mechanisms. Nuclear availability plays an important role in the regulation of NFAT5-dependent gene transcriptions. Under isotonic condition, NFAT5 is localized pan-cellularly as a result of an equilibrium between continuous nuclear import and export mediated by a putative cNLS and a NES respectively (Tong et al., 2006). Upon hypertonic induction, NFAT5 becomes predominantly accumulated in the nucleus, and upregulates NFAT5-dependent gene transcriptions. On the other hand, hypotonicity inactivates NFAT5 by promoting its nuclear export in a NES-independent manner (Tong et al., 2006). A novel nuclear export sequence, which is known as the auxiliary export domain (AED), and which does not bear sequence homology to any known sequences, is crucial for hypotonicity-induced NFAT5 nuclear export(Xu et al., 2008). The corresponding NTRs involved in nuclear import and hypotonicity-mediated nuclear export of NFAT5 have remained elusive. More importantly, how these domains work in a concerted manner to fine-tune the nuclear abundance of NFAT5 in response to changes in tonicity remains unknown.

To understand how NFAT5 activity is regulated by tonicity, we sought to delineate the underlying mechanisms of NFAT5 nucleocytoplasmic trafficking. We found that NFAT5 nuclear import is mediated by a previously uncharacterized NLS, which relies on karyopherin β1 (KPNB1) as the only NTR for the import. We also discovered that hypotonicity-induced nuclear export of NFAT5 is mediated by Exportin-T as the NTR, and requires RUVBL2 as an indispensable chaperone. Our findings have shed lights on the unique nuclear import pathway for cellular adaptation to changes in tonicity, and opened up an opportunity for targeting NFAT5 activity to modulate diverse cellular processes.

## Materials and Methods

### Cell culture and transfections

HeLa cells (American Type Culture Collection, Manassas, VA) were maintained in minimal essential medium supplemented with 10% fetal bovine serum, 1 mM sodium pyruvate, and 2 mM L-glutamine (300 mosmol/kg H2O). For the identification of NFAT5 interacting partners, transfected cells were incubated for 24 h in complete growth medium before switching to hypertonic or hypotonic medium. Hypotonic (260 mosmol/kg H2O) and hypertonic (500 mosmol/kg H2O) media were prepared by supplementing 10% fetal bovine serum, 1 mM sodium pyruvate, 2 mM L-glutamine and NaCl to NaCl-deficient minimal essential medium (Invitrogen) to the desired osmolality. Medium osmolality was measured by the Vapro^®^ vapor pressure osmometer (Wescor). For functional elucidation of NFAT5 interactors using siRNA knockdown, cells were first transfected with the corresponding SMARTpool (Dharmacon) siRNA using DharmaFECT (Dharmacon) for 48 hrs, followed by re-plating cells in three cell culture plates. After 1 day, cells were transfected with the FLAG-NFAT5_132-264_ cDNA overnight, followed by switching to isotonic medium, or switching to hypotonic or hypertonic medium for 30 mins.

### Plasmids, antibodies, siRNAs and inhibitors

FLAG-NFAT5_132-264_ was constructed as described (Xu et al., 2008). Site-directed mutagenesis was conducted using QuikChange site-directed mutagenesis kit (Agilent). FLAG-NFAT5-PEPCK constructs (FLAG-NFAT5_132-264_PEPCK and NFAT5_198-217_PEPCK) were constructed by in-frame insertion of the corresponding NFAT5 and PEPCK into pFLAG-CMV-2 (Sigma-Aldrich) mammalian expression vector (Sigma). FLAG-NFAT5 constructs (FLAG-NFAT5_264-581_, FLAG-NFAT5_132-264_, and FLAG-NFAT5_159-264_) for immunoprecipitation analysis were generated by cloning of the corresponding fragment into pFLAG-CMV-2 vector. NFAT5_174-250_-MiniSOG was constructed by in-frame insertion of NFAT5_171-250_ into pCDNA3.1-MiniSOG (Boassa et al., 2013). GST-NFAT5 constructs (GST-NFAT5_171-250_, GST-NFAT5_171-220_, GST-NFAT5_220-250_, GST-NFAT5_190-250_, GST-NFAT5_190-217_, GST-NFAT5_190-217_) and GST-SV40-NLS were cloned by in-frame insertion of the corresponding NFAT5 fragments and SV40-NLS into the C-terminus of GST of pGEX-4T-1 (GE Healthcare), respectively. Dendra2-Lifeact-7, FLAG-TIP49b (RUVBL2), pET-Ran(Q69L) and AcGFP1-N1 were obtained from (Addgene). Dendra2-RUVBL2 was constructed by in-frame insertion of Dendra2 into the N-terminus of RUVBL2. NFAT5_128-581_-GFP was constructed by in-frame insertion of EGFP to the C-terminus of FLAG-NFAT5_128-581_. siRNA-resistant wild-type (WT) RUVBL2 and siRNA-resistant RUVBL2 ATPase mutant (E300G) were constructed by site-directed mutagenesis of Flag-TIP49b (RUVBL2) using QuikChange site-directed mutagenesis kit (Agilent). His-NFAT5_171-250_-AcGFP, His-SV40_NLS_-AcGFP, and His-KPNB1 were constructed by in-frame insertion of NFAT5_171-250_ and AcGFP, SV40 NLS and AcGFP, and KPNB1 into a modified pET 30a vector, respectively (Pan et al., 2017). His-KPNA5 protein was obtained from LifeSpan BioSciences, Inc. SMARTpool siRNAs for target validations of targets identified by mass spectrometric analysis, against individual karyopherins (IPO4, IPO5, IPO7, IPO8, IPO9, IPO11, IPO13, KPNB1, TNPO1, TNPO3, SNUPN, KPNA1, KPNA2, KPNA3, KPNA4, KPNA5, and KPNA6), and siRNAs against RUVBL1, RUVBL2, JAK1, SLC25A6, MATR3, and IMPDH2, were from Dharmacon. RUVBL2, JAK1, SLC25A6, and IMPDH2 antibodies were obtained from Abcam. ATPase-deficient RUVBL2 was generated by introducing E300G mutation. FLAG antibody (F7425) was from Sigma and was used at 1:1000 (Western blot) and 1:1000 (Immunofluorescence) dilution. NF-90 antibody was from Santa Cruz and was used at 1:1000 dilution. Alpha-Tubulin antibody was from Sigma and was used at 1:1000 dilution. NFAT5 antibody (AAS31388C) obtained from Antibody Verify was used at 1:1000 dilution (Western blot) and NFAT5 antibody (F-9) obtained from Santa cruz was used at 1:100 (Western blot) and 1:50 (Immunofluorescence) dilution. Exportin T (PA5-66095) and Exportin 4 (PA5-65820) were from ThermoFisher and were used at 1:1000 dilution and 1:250 (Immunofluorescence). CB-6644 was obtained from MedChemExpress.

### Photo-oxidation and EM preparation of transfected HeLa cells

Transfected HeLa cells plated on glass bottom culture dishes (MatTek) were fixed in 2.5% glutaraldehyde in 0.1 M cacodylate buffer, pH 7.4, for 1 h on ice, rinsed 5 times in cold cacodylate buffer, and blocked for 30 min with 10 mM KCN, 20 mM aminotriazole, 50 mM glycine, and 0.01% hydrogen peroxide in cacodylate buffer. For MiniSOG photo-oxidation, the transfected cells were identified using a Leica SPE II inverted confocal microscope. Freshly prepared diaminobenzidine (Sigma) in blocking buffer was added to the plate, and HeLa cells were illuminated with 450 – 490 nm light from a xenon lamp for 3–4 min until a light brown reaction product was observed in place of the green fluorescence of MiniSOG. Cells were then removed from the microscope, washed in cold cacodylate buffer, and postfixed in 1% osmium tetroxide for 30 min on ice. After several washes in cold double distilled water, cells were either en bloc stained with 2% aqueous uranyl acetate for 1 h to overnight at 4° C, or directly dehydrated in a cold graded ethanol series (20%, 50%, 70%, 90%, 100%) for 3 min each on ice, then rinsed once in room temperature with 100% ethanol and embedded in Durcupan ACM resin (Electron Microscopy Sciences). Sections were cut with a diamond knife at a thickness of 70 –90 nm for thin sections and 250 nm for thick sections for electron tomography. Thin sections were examined using a JEOL 1200 EX operated at 80 kV.

### Electron tomography

Sections were coated with carbon on both sides, and colloidal gold particles (5 and 10 nm diameter) were deposited on each side to serve as fiducial markers. For reconstruction, double or triple tilt series of images were recorded at regular tilt (angular increments of 2° from -60° to +60° increments) with a JEOL 4000EX intermediate high-voltage electron microscope operated at 400 kV. The specimens were irradiated before initiating a tilt series to limit anisotropic specimen thinning during image collection. Tilt series were recorded using a 4k X 4k custom high resolution slow-scan CCD camera system delivering 25% contrast at Nyquist. Fine alignment of projections and 3D reconstruction were performed using the TxBR reconstruction package (Lawrence et al., 2006). This high-resolution tomography reconstruction software was used in conjunction with the IMOD package, where reconstructed image volumes were viewed and objects of interest traced manually and reported as 3D models

### Immunofluorescence and confocal microscopy

Cells were washed three times with PBS and fixed with 4% w/v paraformaldehyde for 15 min at 4 °C, followed by permeabilization with absolute methanol for 2 min at room temperature. Primary antibodies against FLAG (Sigma) were used. Alexa 488 anti-rabbit antibody (Invitrogen) was used as secondary antibodies. To visualize the nuclei, cells were stained with 4,6-diamidino-2-phenylindole (DAPI) (Sigma). Images were viewed either with a Zeiss Axiovert 200 M fluorescence microscope or Leica TCS SP8 MP confocal microscope system. Quantification of fluorescence signals in different subcellular compartments was conducted to our previously described method (Xu et al., 2008; Tong et al., 2006).

### Immunoaffinity purification, multidimensional Protein Identification Technology (MudPIT), mass spectrometry, and data base searching

HeLa cells were transfected with FLAG-NFAT5132-581. Twenty-four hours after transfection, cells were pre-treated with hypotonic medium (260 mosmol/kg H2O) for 60 mins and switched to hypertonic medium (450 msomol/kg H2O) for 30 mins, and then lysed with lysis buffer (50 mM Tris-Cl, 150 mM NaCl, 1mM EDTA, pH 7.5, 1% Triton X-100, cocktail protease inhibitors and phosphatase inhibitors). The recombinant protein was purified by affinity chromatography using anti-FLAG affinity resin (Sigma). The purified protein was precipitated by 20% trichloroacetic acid. The precipitate was washed twice with acetone, dried, and resuspended in 8 M urea and 100 mM Tris-HCl, pH 8.5. The solubilized protein was reduced by the addition of Tris(2-carboxyethyl)phosphine to 5 mM, followed by the carboxyamidomethylation of cysteines using 10 mM iodoacetamide. The concentration of urea was then diluted 2-fold (to 4 M) by the addition of an equal volume of 100 mM Tris-HCl, pH 8.5. Sequencing grade endoproteinase Lys-C (Roche Diagnostics) and modified trypsin (Promega) were added at ∼1:50 enzyme to substrate ratio (w/w) and incubated at 37 °C for 4 and 12–16 h, respectively. The resulting peptides were extracted with 5% formic acid and redissolved into buffer A (5% acetonitrile with 0.1% formic acid). Data-dependent tandem MS analysis was performed with a LTQ-Orbitrap mass spectrometer (Thermo Fisher Scientific). Protein identification was done with Integrated Proteomics Pipeline (IP2, Integrated Proteomics Applications, Inc. San Diego, CA. http://www.integratedproteomics.com), using ProLuCID algorithm to conduct database search (Eng et al., 1994; Xu et al., 2015), and DTASelect2 to filter result (Cociorva et al., 2007; Tabb et al., 2002). Tandem mass spectra were extracted into ms1 and ms2 files from raw files using RawExtract 1.9.9 (http://fields.scripps.edu/downloads.php) and then were searched against EBI-IPI human protein database (version 3_71, released on 03-24-2010) (McDonald et al., 2004). To estimate peptide probabilities and false-discovery rates accurately, we used a reverse decoy database containing the reversed sequences of all the proteins appended to the target database (DuMond et al., 2016). All searches were parallelized and performed on Intel Xeon 80 processor cluster under the Linux operating system. The peptide mass search tolerance was set to 10 ppm for spectra acquired on the LTQ-Orbitrap instrument. The mass of the amino acid cysteine was statically modified by +57.02146 Dalton, to take into account the carboxyamidomethylation of the samples and 2 peptides per protein and at least one trypitic terminus were required for each peptide identification. The ProLuCID search results were assembled and filtered using the DTASelect program (version 2.0) with false discovery rate (FDR) of 0.05; under such filtering conditions, the estimated false discovery rate was below ∼1% at the protein level in all analyses.

### Live cell microscopy and time-lapse imaging

Live cell microscopy and time-lapse imaging were conducted as described previously (Xu et al., 2008). HeLa cells transiently expressed NFAT5_128-581_-EGFP fusion proteins were viewed with an inverted Zeiss fluorescence microscope. A heated chamber perfused with CO2 was used to incubate the cells at 37 °C. After pre-treatment of cycloheximde for 1 hr (5ug/ml; Tong et al. 2006), the original growth medium was removed and replaced with a pre-warmed hypotonic growth medium. Images were taken at 30-min intervals and analyzed using NIH ImageJ software (http://imagej.nih.gov/ij/). Quantification of signal intensity was conducted according to our previously described method (Ramírez et al., 2009). A circular ROI (25 pixels in diameter) was selected within the nucleus and cytoplasm of each cell for the quantification of signal intensity. The average signal intensity was then subtracted with the average background signal intensity.

### Photoconversion of Dendra2 and analysis of nuclear export of Dendra2 -

RUVBL2 – Photoconversion of Dendra2-RUVBL2 from green to red fluorescence was achieved by exposing the cells to blue light according to a described method (Gurskaya et al., 2006). Cell nucleus was defined and exposed to 405 nm laser light for 4-5 times 200ms bursts at ∼5% 405 nm laser power (TCS SP8 MP, Leica). After photoconversion of Dendra2-RUVBL2, the original growth medium was removed and replaced with a pre-warmed hypotonic growth medium. Images were taken at 10-min intervals. Post-acquisition image analyses were performed using NIH ImageJ software (http://imagej.nih.gov/ij/). A circular ROI (12 pixels in diameter) was selected within the nucleus and cytoplasm of each cell for the quantification of signal intensity. The average signal intensity was then subtracted with the average background signal intensity.

### Protein Extraction, western blotting, and immunoprecipitation

Cell protein lysate was obtained from cell culture using lysis buffer (50 mM Tris-HCl pH7.8, 100 mM NaCl, 1 mM EDTA, 1% Triton X-100, 1 mM phenylmethylsolfonyl fluoride, and 10 mM dithiothreitol). Cytoplasmic and nuclear extracts were prepared using Nuclear and Cytoplasmic Extraction reagents (ThermoFisher Scientific). SDS-PAGE electrophoresis, Western Blotting, and immunoprecipitation were conducted as described (Chen et al., 2013).

### Protein expressions and purifications

Full-length KPNB1 and various NFAT5 fragments were cloned into expression vectors with GST, 6xHis or Trx-6xHis-3C fusion tag, and transformed into E. coli BL21 (DE3) bacteria (Invitrogen). Bacteria were harvested, lysed in lysis buffer (PBS with 5% glycerol, 100 mM MgCl_2_, 1 mM PMSF, 1 mM DTT, 1 mg/ml lysozyme, 0.2 units/ml DNase I, and Complete protease inhibitors), and centrifuged to remove debris. GST fusion proteins were purified from lysates with glutathione-Sepharose 4 Beads (GE Healthcare). His-tag fusion proteins were purified from lysates using HisTrap HP column (GE Healthcare). The size and purity of the protein preparations were verified by SDS-PAGE and Coomassie Blue staining. The Trx-6xHis-3C tag was removed by protease 3C cleavage and the untagged protein was further purified by size-exclusion chromatography (Superdex 75, GE Healthcare).

### Isothermal Titration Calorimetry

Isothermal Titration Calorimetry (ITC) was performed using the MicroCal^TM^ PEAQ-ITC (Malvern Panalytical Ltd). NFAT5_151-216_, NFAT5_189-216_ and full length KPNB1 were dialyzed into 50 mM Tris, pH 8.0, and 150 mM NaCl. NFAT5_171-253_ and full length KPNB1 were dialyzed into 10 mM Na_2_HPO_4_·7H_2_O, 1.8 mM KH_2_PO_4_, 2.7mM KCl, pH 7.4 and 150 mM NaCl. 150 μl of NFAT5_151-216_, NFAT5_189-216_ proteins at 1.5-1.75mM were respectively titrated into the sample cell loaded with 400 μl of full length KPNB1 at 50 μM. 150 μl of NFAT5_171-253_ proteins at 600 μM were titrated into the sample cell containing 400μl of 20 μM KPNB1. Typically, titrations consisted of 19 injections of 2 μl, with 150-s equilibration between injections. The data were analyzed using MicroCal PEAQ-ITC Analysis Software.

### *In-Vitro* pulldown assay

GST-fusion proteins, His-KPNB1 and Ni-NTA agarose were mixed and incubated at 4°C for with shaking overnight. The mixtures were centrifuged at 5000 rpm, 4°C for 2 minutes. The flow through was discarded. Agarose was washed three times with buffer (20 mM sodium phosphate, 100 mM sodium chloride, 40 mM imidazole, 10% glycerol). Proteins were eluted from agarose by boiling SDS-containing buffer for 10 minutes. After spinning down the debris, the supernatant was analyzed by SDS-PAGE and visualized by Coomassie Blue staining.

### Reconstitution of nuclear import in permeabilized cells

The assay was conducted as described (Cassany and Gerace, 2009). In brief, HeLa cells were grown in 10-well slides (Millipore). Cells were washed three times with cold transport buffer (20 mM Hepes, pH7.3, 110 mM potassium acetate, 2 mM magnesium acetate, 1 mM EGTA, 2mM DTT, 1 mM PMSF, protease inhibitor) on ice, followed by permeabilization with 50 μg/mL digitonin at room temperature for 5 minutes. Permeabilized cells were washed three times with cold transport buffer. Reaction was carried out in the presence of ATP-regenerating system (1 mM ATP, 1 mg/mL CP, and 15 U/mL CPK), 0.1 mM GTP, recombinant proteins (His-KPNB1, His-KPNA5, NTF2, RAN, His-NFAT5-AcGFP) and various supplements whenever necessary (0.8 mg/mL WGA, 2 mg/mL cytosol preparation, 0.2 mM GTPγS). After incubation, the cells were washed three times with cold transport buffer on ice and fixed with 4 % paraformaldehyde on ice for 10 minutes, followed by permeabilization with 100% methanol for 5 minutes at room temperature. Nuclei were stained using 4’, 6-Diamidine-2’-phenylindole dihydrochloride (DAPI) for 5 minutes. After washing, cells were mounted on a glass slide by FluorSave™ Reagent (Millipore). The GFP and DAPI signals were observed using confocal microscope (Leica TCS SP8).

### Statistical analysis

Statistical analysis was conducted using GraphPad Prism software. All data were expressed as mean ± SEM of triplicate experiments. Statistical significance was determined by unpaired t-test (two-tailed) for two groups or one-way ANOVA or two-way ANOVA for three or more groups comparison, followed by Bonferroni’s multiple comparison test as post-test. Significance is shown in the respective figures. p< 0.05 was considered statistically significant.

## Results

### An extensive NLS is required for nuclear import of NFAT5

Our earlier studies showed that FLAG-OREBP1-581Δ1-131 (renamed hereafter as FLAG-NFAT5_132-581_) (Xu et al., 2008; Tong et al., 2006) (Fig. 1A), a truncated form of NFAT5 devoid of both the N-terminal NES and C-terminal transactivation domain, faithfully recapitulated nucleocytoplasmic trafficking properties of endogenous NFAT5 in response to changes in extracellular tonicity. This truncated recombinant NFAT5 contains an AED (a.a.132-156) indispensable for nuclear export under hypotonicity; a putative cNLS (a.a. 199-216, here we called it NFAT5-cNLS) as suggested by Motif Scan (http://myhits.isb-sib.ch/cgi-bin/motif_scan); and a Rel-homology DNA-binding domain (RHD) (a.a. 264 – 581)(Tong et al., 2006; Xu et al., 2008). However, it does not contain the NES (a.a. 8-15) that is responsible for nucleocytoplasmic shuttling of NFAT5 under isotonic condition(Tong et al., 2006; Cheung and Ko, 2013). Therefore, FLAG-NFAT5_132-581_ is constitutively localized to the nucleus under isotonic condition, and the AED drives nuclear export activity of this recombinant protein under hypotonicity(Tong et al., 2006). We generated deletion mutants to decipher the minimal protein domains required for NFAT5 nucleocytoplasmic trafficking (Fig. 1A). First, we elucidated if RHD is dispensable by substituting it with cytoplasmic phosphoenolpyruvate carboxykinase (PEPCK-C). PEPCK-C is an exclusively cytoplasmic protein used as a reporter for the characterization of NLS activity(Theodore et al., 2008). The apparent molecular weight of the new fusion protein (FLAG-NFAT5_132-264_PEPCK) is 83 kDa (Supplementary Fig. 1A), which precludes a passive nuclear transport mechanism. Both FLAG-NFAT5_132-581_ and FLAG-NFAT5_132-264_PEPCK were predominantly localized to cytoplasm and nucleus when cells were subjected to hypotonic and hypertonic challenge respectively (Fig. 1B). The replacement of RHD by PEPCK resulted in a modest reduction in nuclear localization of the recombinant protein under both isotonic and hypertonic condition, suggesting that RHD has a minimal impact on NFAT5 nuclear import. However, a PEPCK fusion protein consisted of the putative NFAT5-cNLS and PEPCK (FLAG-NFAT5198-217PEPCK) (Fig. 1A) failed to enter nucleus regardless of extracellular tonicity (Fig. 1B). As a control, FLAG-PEPCK was exclusively localized to the cytoplasm under all tonicities examined. Our findings suggested that amino acid residues 132-264 of NFAT5 confer tonicity-sensitive nuclear import and export activity to a heterologous protein, whereas the NFAT5-cNLS *per se* is an inactive nuclear import signal. Nevertheless, alanine substitution of the three core residues (202RKR204) in NFAT5-cNLS resulted in complete ablation of nuclear import of NFAT5(Tong et al., 2006), suggesting that NFAT5-cNLS is essential but insufficient to confer significant nuclear import activity. To further delineate the minimal essential sequence required for nuclear import activity, we conducted nested deletion of FLAG-NFAT5_132-264_PEPCK from the N- and C-terminus respectively, and determined nucleocytoplasmic localization of the truncated proteins under different tonicities. As shown in Fig. 1C, consistent with our earlier findings that the AED is important for nuclear export(Tong et al., 2006), AED-deficient fusion protein (FLAG-NFAT5_159-264_PEPCK) became constitutively localized to the nucleus under different tonicities. Deletion mapping analysis suggested that a.a. residues 159 to 173 from the amino terminus (FLAG-NFAT5_174-264_PEPCK) were not required for nuclear trafficking of the fusion protein. Nevertheless, nuclear localization of the fusion protein was profoundly inhibited when a.a. residues 174 to 188 (FLAG-NFAT5_189-264_PEPCK) were removed, and completely abolished when a.a. residues 189 to 197 (FLAG-NFAT5_198-264_PEPCK) were deleted. On the other hand, a.a. residues from 264 to 251 from the C-terminus (FLAG-NFAT5_174-250_PEPCK) were not required for nuclear localization of the fusion protein, but deletion of a. a. residues from 250 to 241 (FLAG-NFAT5_174-240_PEPCK) impaired nuclear localization significantly. Furthermore, deletion of a. a. residues from 240-231 (FLAG-NFAT5_174-230_PEPCK) resulted in exclusive cytoplasmic localization of the fusion protein. Together, these data suggested that the presuming NFAT5-cNLS (Fig. 1D) does not serve as a functional NLS. A minimal protein domain comprised of 52 amino acids (a.a. residues 189-240) is required for directing nuclear import of NFAT5. However, a domain comprised of 77 amino acids (a. a. residues 174 to 250) exhibited full nuclear import activity (Fig. 1D). We named this NLS the NFAT5-NLS (Fig. 1D).

**Figure 1.**
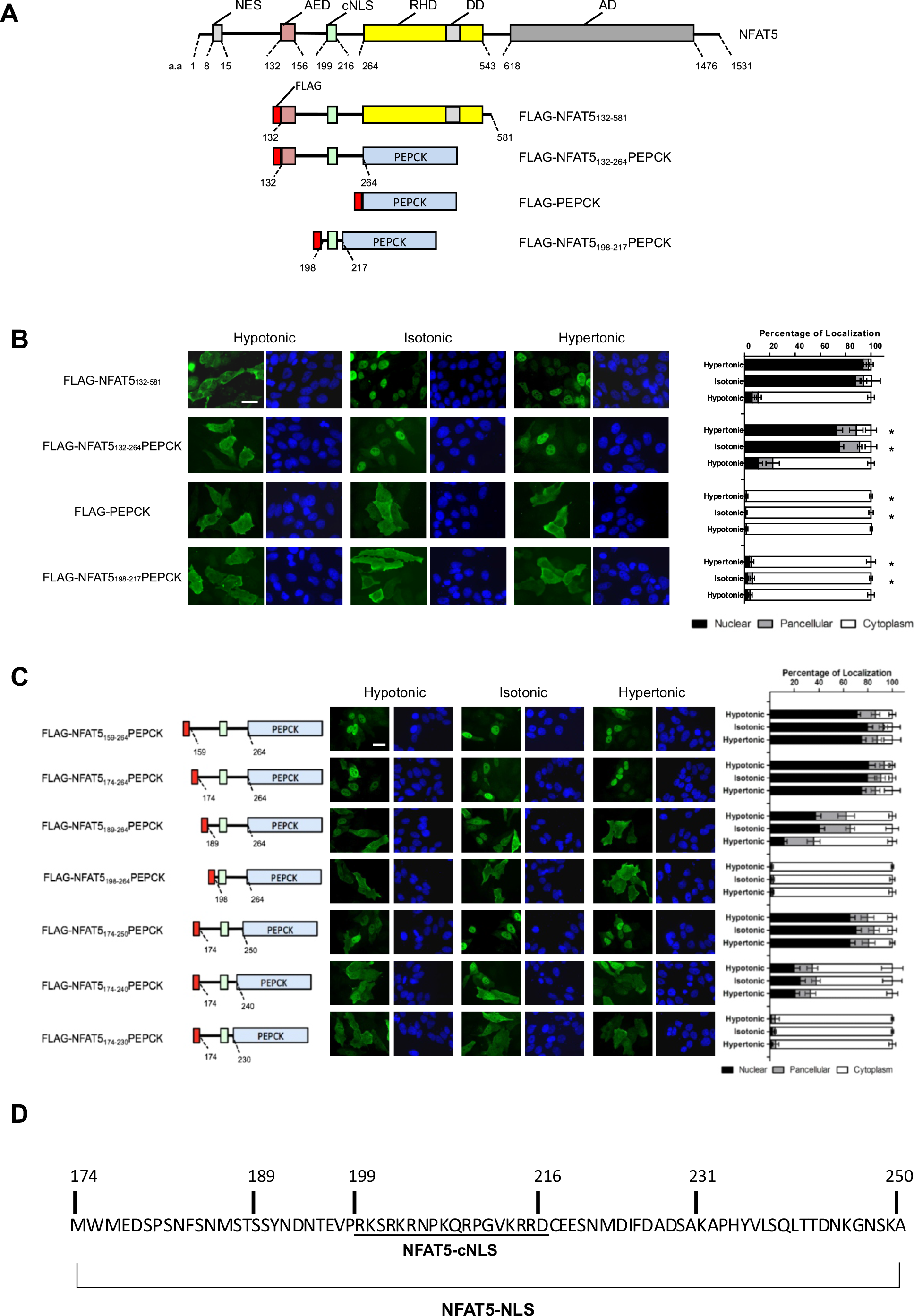
**A)** Schematic illustration of NFAT5, FLAG-NFAT5_132-581_ and FLAG-NFAT5-PEPCK fusion constructs. NES, canonical nuclear export signal; AED, axillary export domain; cNLS, classical nuclear localization signal; RHD, Rel-homology domain; DD, dimerization domain; AD, putative transcriptional activation domains. **B)** Left, representative fluorescence images, scale bar is 30 µm. Right, quantitative analysis of subcellular localization of fluorescence signal in HeLa cells expressing recombinant NFAT5 and FLAG-NFAT5-PEPCK mutants, treated with hypotonic, isotonic or hypertonic medium for 90 mins. Cells were stained with FLAG antibody and FITC-labeled secondary antibody, counterstained with DAPI. **C)** Mapping of minimal essential NLS using FLAG-NFAT5-PEPCK deletion mutants. Left schematic illustration of the mutants; Middle, representative fluorescence images of cells expressing recombinant FLAG-NFAT5-PEPCK mutants, scale bar is 30 µm; Right, quantitative analysis of subcellular localization of fluorescence signal in HeLa cells expressing indicated plasmids. Cells were switched to hypotonic, isotonic or hypertonic medium for 90 mins before fixation. Cells were stained with FLAG antibody and FITC-labeled secondary antibody, counterstained with DAPI. In B and C, at least 100 cells were scored in each condition. Data are presented as mean ± SEM of three independent experiments. *, p < 0.0001 by one-way ANOVA with Bonferroni’s multiple comparison test as post-test. **D)** Amino acid sequences of the NFAT5-NLS required for NFAT5 nuclear import. cNLS, classical NLS.

### Nucleocytoplasmic trafficking of NFAT5 is mediated through the nuclear pore

To visualize nucleocytoplasmic trafficking of NFAT5 in response to changes in extracellular tonicity *in situ*, we conducted Correlated Light and Electron Microscopy (CLEM) analysis (Ellisman et al., 2012). CLEM takes advantage of the Mini Single Oxygen Generator (MiniSOG), a small (106 a.a.) fluorescent flavoprotein that generates reactive oxygen species upon illumination of blue light(Ludwig et al., 2013). Local production of reactive oxygen species in glutaraldehyde-stabilized cells then photooxidize diaminobenzidine (DAB) into an osmiophilic electron-dense polymer, which allows proteins to be visualized at high spatial resolution and contrast under electron microscopy (EM), with good preservation of ultrastructures(Ludwig et al., 2013; Ou et al., 2012). We generated NFAT5-MiniSOG fusion gene (NFAT5_174-250_-MiniSOG). The fusion protein, which contains both AED and NFAT5-NLS, was subjected to cytoplasmic and nuclear localization treatment with hypotonicity and hypertonicity respectively (Fig. 2A). Photooxidation of glutaraldehyde-fixed NFAT5_174-250_-MiniSOG expressing cells in the presence of DAB resulted in differential deposition of a brown reaction products. Electron micrographs of plastic embedded and osmium-stained sections revealed that electron-dense staining was primary localized to the cytoplasm and nucleus in response to hypotonic and hypertonic treatment respectively (Fig. 2B). Under hypotonic condition, at higher magnification, ring-shaped staining at nuclear rim was observed, consistent with the structure of nuclear pores (yellow arrows in Fig. 2B, right panel a’), suggesting that the fusion proteins were captured at the nuclear pore during nuclear export. Under hypertonic condition, ring-shaped staining at nuclear rim was also observed, which is consistent with the structure of nuclear pores (yellow arrows in Fig. 2B, right panel b’). Interestingly, imaging by electron tomography revealed a DAB-positive patchy staining within the nucleus. which distributed exclusively in regions outside of the most condensed chromatin (black arrows in Fig. 2C), consistent with the notion that active transcription occurs at the fringes of chromosome territories(Steensel and Furlong, 2019). Three-dimensional electron tomographic analysis of hypertonicity-treated cells further confirmed increased staining density at the nuclear pore (yellow arrows in Fig. 2B). Together these data provide direct evidence that NFAT5 distributes in the nucleus in areas with less condensed chromatin and undergoes nucleocytoplasmic trafficking through nuclear pore complex.

**Figure 2.**
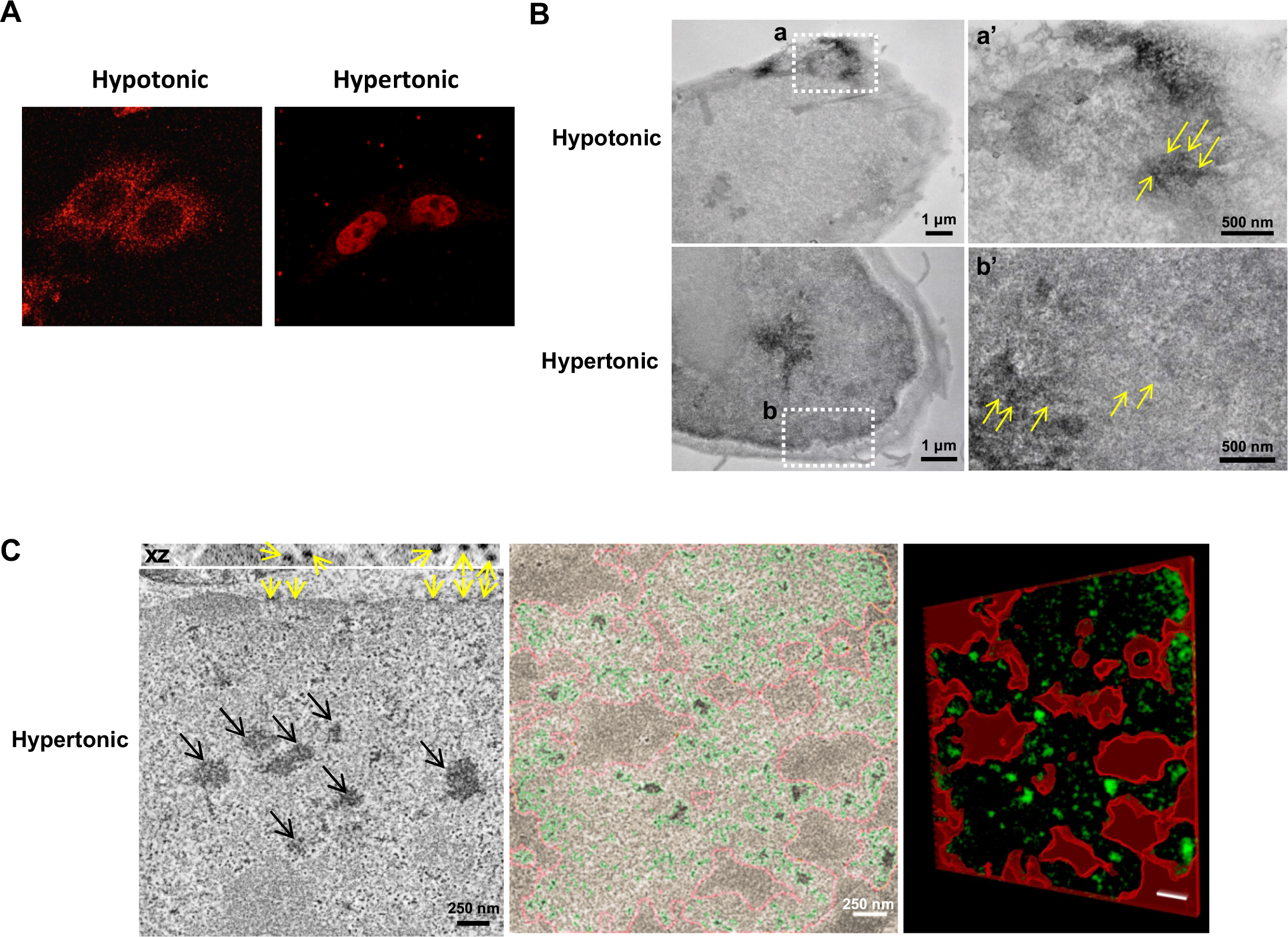
**A)** HeLa cells transfected with NFAT5_174-250_-MiniSOG. Fluorescence signals were visualized by confocal microscopy (100X); **B)** Spatial distribution of NFAT5_174-250_-MiniSOG following different tonicities by electron microscopy. Following photo-induced oxidation of DAB, NFAT5_174-250_-MiniSOG-expressing HeLa cells were processed and visualized by electron microscopy. Under hypotonic condition, the DAB-positive staining was clearly observed in the cytoplasm. The high magnification image (a’) corresponds to the same area indicated by the white square (a). Yellow arrows indicate staining associated with nuclear pores. In contrast, under hypertonic condition, the intense DAB-positive staining was observed in the nucleus. The high magnification image (b’) corresponds to the same area indicated by the white square (b). Yellow arrows indicate staining associated with nuclear pores. **C)** Three-dimensional electron tomographic analysis of hypertonicity-treated HeLa cells showed the pattern of staining within the nucleus exclusively associated with regions of less condensed chromatin (as indicated by the black arrows, left image). Yellow arrows indicate the staining at the nuclear pores, particularly visible in the xz plane. Segmentation of a tomogram based on thresholding highlights the NFAT5 staining (in green) exclusively outside of the areas of more condensed chromatin (delimited in red, middle and left image).

### KPNB1 is the nuclear import receptor for NFAT5

We sought to determine the NTR responsible for directing NFAT5 nuclear import. To systematically evaluate the involvement of Kapβ in this process, we conducted siRNA knockdown to deplete Kapβs that are known to mediate nuclear import via recognizing distinct classes of NLS on import cargoes(Soniat and Chook, 2015), followed by determination of subcellular localization of FLAG-NFAT5_132-264_PEPCK. Among others, only depletion of KPNB1 (Impβ1), but not TNPO1 (Kapβ2), IPO4 (Imp4), IPO5 (Imp5), IPO7 (Imp7), IPO8 (Imp8), IPO9 (Imp9), IPO11 (Imp11), IPO13 (Imp13), or TNPO3 (Trn-SR), markedly inhibited nuclear localization of FLAG-NFAT5_132-264_PEPCK under both isotonic and hypertonic conditions (Fig. 3A and Supplementary Fig. 1B). Nuclear abundance of NFAT5 was diminished significantly in KPNB1-depleted cells (Fig. 3B). In addition, co-immunoprecipitation of FLAG-NFAT5_132-581_ resulted in the presence of endogenous KPNB1 in the immunocomplex (Fig. 3C), suggesting that KPNB1 is associated with NFAT5. KPNB1 was known to heterodimerize with nuclear import adaptor karyopherin-alpha (Kapα) for nuclear transport of cNLS-containing cargo(Marfori et al., 2011). siRNA knockdown was used to determine if any members of the Kapα family, including KPNA1 (Impα1), KPNA2 (Impα2), KPNA3 (Impα3), KPNA4 (Impα4), KPNA5 (Impα5), KPNA6 (Impα6), and KPNA7 (Impα7), or SNUPN (Snurportin 1), a novel nuclear import adaptor of KPNB1(Huber et al., 1998), are involved in the NFAT5 nuclear import. Nuclear localization of FLAG-NFAT5_132-264_PEPCK was not significantly altered by the knockdown of any of these Kapα members (Fig. 3D), suggesting that KPNB1 mediates nuclear import of NFAT5 in the absence of nuclear import adaptors from the Kapα family.

**Figure 3.**
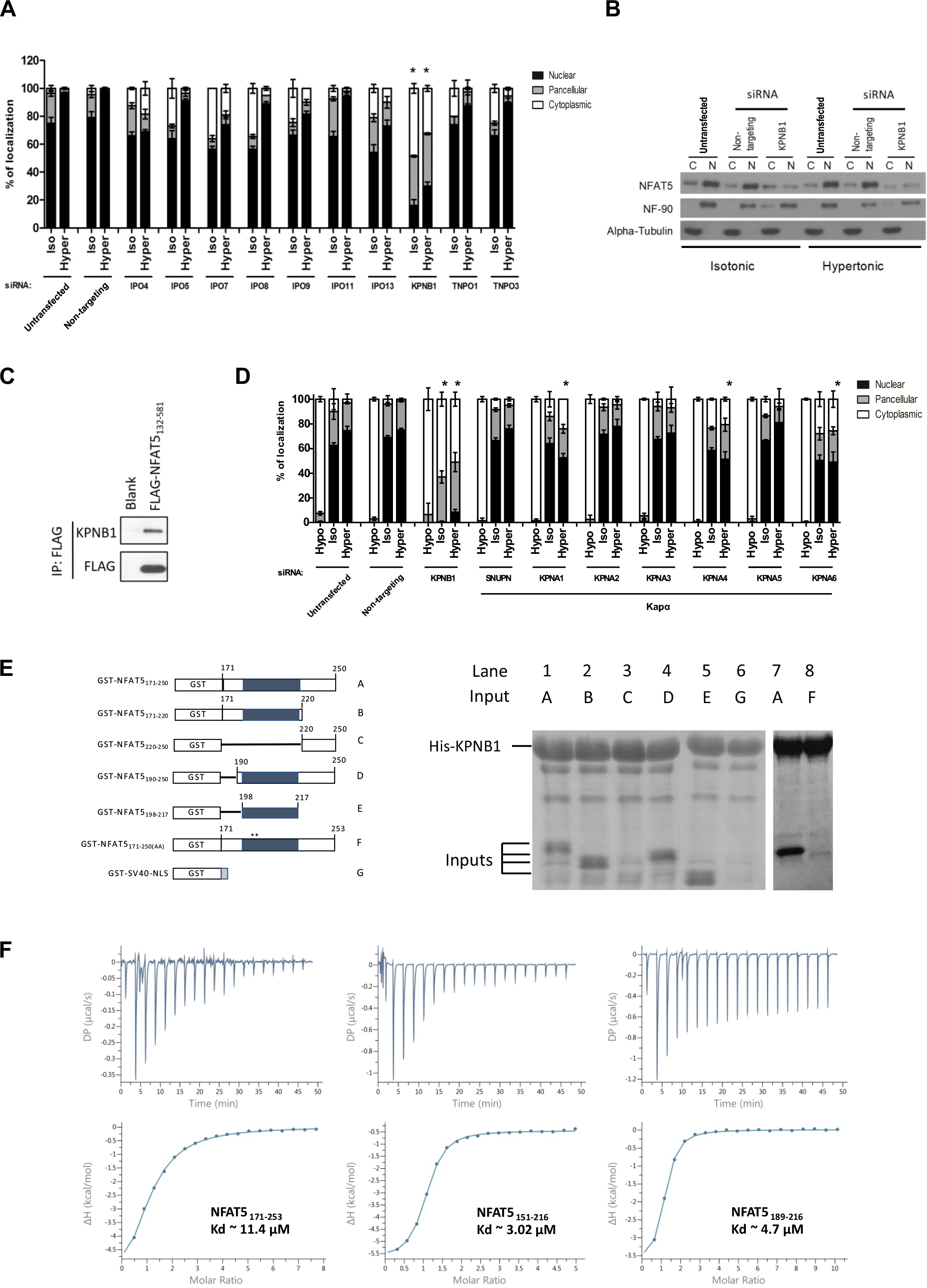
**A**) Quantitative analysis of subcellular localization of fluorescence signal in HeLa cells transfected with FLAG-NFAT5_132-264_PEPCK and the indicated siRNA from the Kapβ family, followed by treatment with hypotonic, isotonic or hypertonic medium for 90 mins. Cells were stained with FLAG antibody and FITC-labeled secondary antibody, counterstained with DAPI. **B)** Western blotting analysis showing cytoplasmic (C) and nuclear (N) distribution of endogenous NFAT5 in HeLa cells transfected with siKPNB1 under different extracellular tonicities. NF-90 and alpha-tubulin antibodies were used as nuclear and cytoplasmic marker, respectively. **C**) Co-immunoprecipitation analysis of FLAG-NFAT5_132-581_ and KPNA1. Immunoprecipitation was carried out using anti-FLAG affinity resin and immunocomplexes were subjected to Western blot analysis using the KPNB1 and FLAG antibodies, respectively. **D**) Quantitative analysis of subcellular localization of fluorescence signal in HeLa cells expressing FLAG-NFAT5_132-581_ and the indicated siRNA of members in the Kapα family. Cells were switched to hypotonic, isotonic or hypertonic medium for 90 mins before fixation. Cells were stained with FLAG antibody and FITC-labeled secondary antibody, counterstained with DAPI. In A and D, at least 100 cells were scored in each condition. Data are presented as mean ± SEM of three independent experiments. *, represents significant reduction in nuclear signal compared with the untransfected cells under the same tonicity; p<0.0001 by one way ANOVA with Bonferroni’s multiple comparison test as post-test. **E)** Left, schematic representation of different GST-NFAT5 and GST-SV40-NLS fusion proteins. Right, i*n vitro* pull-down analysis using His-KPNB1. For lanes 1-8, His-KPNB1 was immobilized on Ni-NTA agarose followed by the addition of the indicated GST-NFAT5 fusion protein. After washing, proteins were eluted with excess imidazole. Proteins were analyzed by SDS-PAGE followed by Coomassie Blue staining. **F)** Isothermal Titration Calorimetry (ITC) profiles to measure the binding affinity of NFAT5 to KPNB1. Different constructs of NFAT5 putative NLS region, including residues 171-253, 151-216 and 189-216 that showed similar binding affinities (Kd).

To confirm if KPNB1 is the sole NTR for NFAT5 nuclear import, we conducted *in vitro* pull-down assay to test whether recombinant NFAT5-NLS proteins (Fig. 3E, left, and Supplementary Fig. 1C) and KPNB1 interact directly. Agarose beads coupled to His-KPNB1 efficiently pulled down GST-NFAT5_171-250_, but not GST control (lane 1 vs lane 6). However, truncation of GST-NFAT5_171-250_ from either terminus did not appreciably weaken its interaction with KPNB1 (lane 2 and 4 vs lane 1). Intriguingly, despite the failure of NFAT5-cNLS to confer nuclear import activity to PEPCK, GST-NFAT5198-217, which contains NFAT5-cNLS only, interacts significantly with His-KPNB1 at a level comparable to GST-NFAT5_171-250_ (lane 5 vs lane 1). However, their interaction was completely abolished when the NFAT5-cNLS (a.a. 199-216) was removed (lane 3), or when the core basic amino-acids (R202 and K203) were mutated to alanine (lane 8), a result that was consistent with the notion that these residues are important for nuclear import activity. Together, these data suggested that although NFAT5-NLS is required for nuclear import activity, and that the NFAT5-cNLS is sufficient for establishing interaction with KPNB1. To further understand this interaction, we conducted isothermal titration calorimetry (ITC) experiments to measure the binding affinity of different NFAT5 fragments to full-length Kpnb1. We found that NFAT5171-253 and NFAT5_151-216_ bind to KPNB1 with almost the same binding affinity as NFAT5_189-216_ that only comprises the NFAT5-cNLS (*K_d_* ∼ 1.2 μM and 1.8 μM vs. 1.8 μM) (Fig. 3F). These ITC measurements corroborated with our pull-down assays and further showed that different NLS fragments interact with KPNB1 at similar affinity.

### *In vitro* reconstitution of KPNB1-mediated nuclear import

As there is a discrepancy between the amino acid residues of NFAT5 required for effective KPNB1 interaction *in vitro*, and effective nuclear import *in vivo*, *in vitro* nuclear transport assay was conducted using digitonin-permeabilized HeLa cells to further discern the underlying mechanism. Recombinant His-tagged NFAT5-NLS fused to monomeric green fluorescent protein (AcGFP) (His-NFAT5171-250-AcGFP), His-tagged SV40 Large-T antigen NLS fused to AcGFP (His-SV40_NLS_-AcGFP), and His-tagged AcGFP (His-AcGFP) were bacterially expressed and purified (Supplementary Fig. 1D), and their nuclear import activities were evaluated. Both His-NFAT5171-250-AcGFP and His-SV40_NLS_-AcGFP underwent nuclear import dependent on the presence of cytosolic extracts, ATP-regeneration mixture, and RanGTP, whereas the import was abolished in the presence of nuclear pore inhibitor wheat germ agglutinin (WGA), high concentration of GTP, or nonhydrolyzable GTP analog (GTPγS) (Supplementary Fig 3B). His-AcGFP, which does not contain any NLS, failed to enter nucleus (Supplementary Fig. 1E). These findings suggested that, similar to His-SV40_NLS_-AcGFP, His-NFAT5171-250-AcGFP enters nucleus in an ATP- and GTP-dependent manner mediated by cytosolic factors, presumably via the NTRs.

To further delineate molecular determinants for His-NFAT5171-250-AcGFP nuclear import, we conducted *in vitro* nuclear transport assay by NTRs and defined factors. In the absence of exogenous factors, His-NFAT5171-250-AcGFP or His-SV40_NLS_-AcGFP did not undergo nuclear import (Fig. 4A, lane 1). Nuclear translocation of both recombinant proteins was evidenced in the presence of RanGTP, nuclear transport factor 2 (NTF2), ATP-regenerating mixture, KPNA5 and KPNB1 (Fig. 4A, lane 2), but was abolished by the addition of WGA (Fig. 4A, lane 5). In agreement with the notion that nuclear import of cNLS requires ternary complex formation with importin-α and - β, nuclear accumulation of His-SV40-AcGFP was abolished in the absence of KPNA5 (Fig. 4A, lane 4) or KPNB1 (Fig. 3E, lane 5). In contrast, nuclear import of His-NFAT5171-250-AcGFP was abolished in the absence of KPNB1 (Fig. 4A, lane 5) but not KPNA5 (Fig. 4A, lane 4). Therefore, the mechanism of NFAT5 nuclear import is mediated by a KPNB1-dependent and Kapα-independent mechanism, which is distinctive from the classical SV40 T-antigen nuclear import pathway.

**Figure 4.**
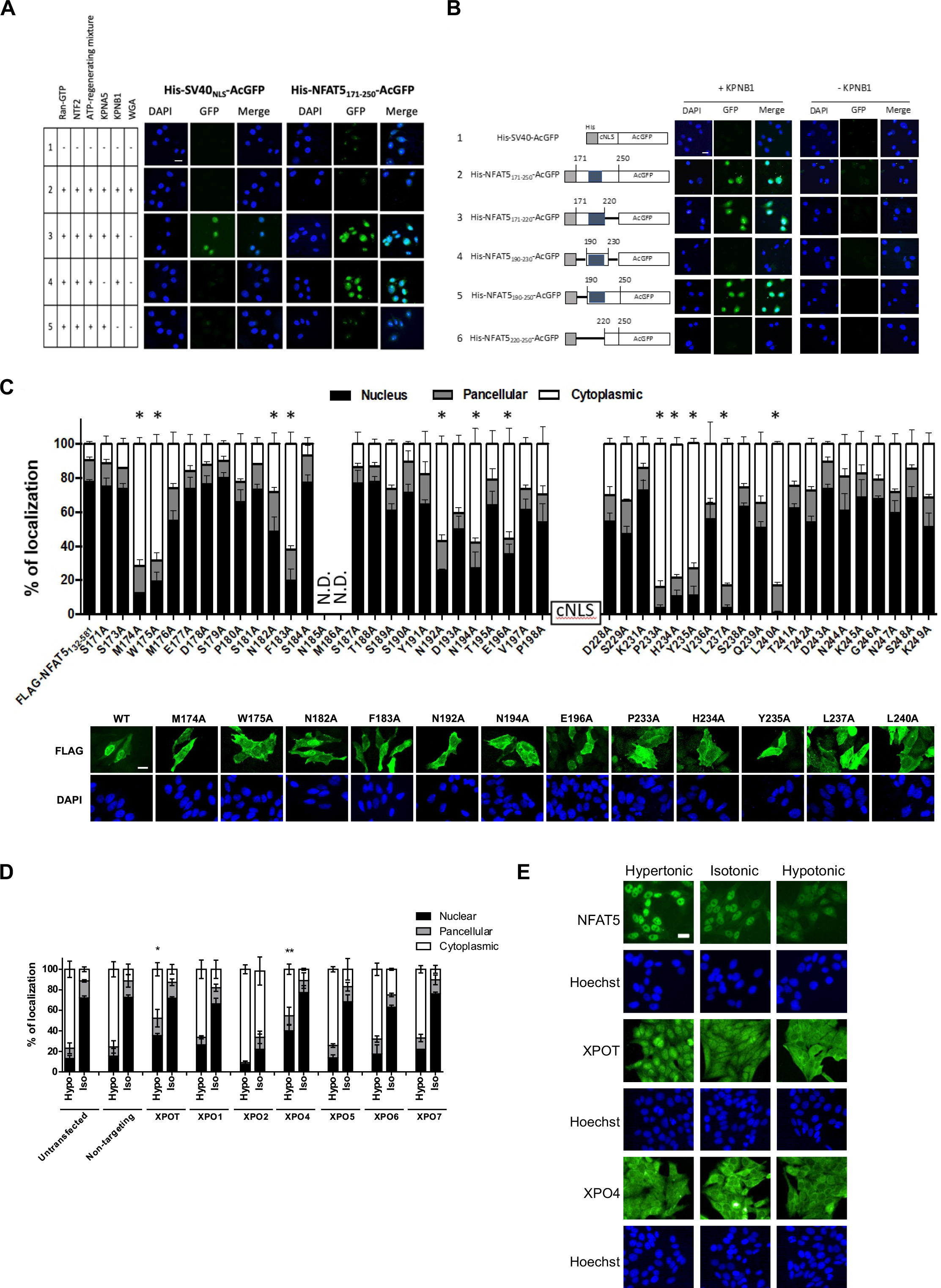
**A)** *In vitro* nuclear import assay of His-SV40_NLS_-AcGFP and His-NFAT5171-250-AcGFP was carried out using digitonin-permeabilized HeLa cells supplemented with individual nuclear transport factors as specified. Transport assay was carried out for 30 min at 37°C. Cells were then fixed with paraformaldehyde, stained with DAPI, and GFP fluorescence images were taken using confocal microscopy. Scale bar is 30 µm. **B)** Role of KPNB1 in nuclear import of His-NFAT5-AcGFP. Left, schematic representation of different His-NFAT5-AcGFP reporter constructs. Right, *in vitro* nuclear import assay of His-SV40-AcGFP and His-NFAT5-AcGFP was carried out using digitonin-permeabilized HeLa cells supplemented with RanGTP, NTF2, and ATP-regenerating mixture, in the presence or absence of KPNB1. Transport assay was carried out for 30 min at 37°C. Cells were then fixed with paraformaldehyde, stained with DAPI, and images were taken using confocal microscopy. Scale bar is 30 µm. **C)** Top, quantitative analysis of subcellular localization of fluorescence signal in HeLa cells expressing alanine-substituted FLAG-NFAT5 mutants. 24 hours after transfection of FLAG-NFAT5_132-581_ carrying the indicated mutation under isotonic condition, cells were fixed with paraformaldehyde, stained with DAPI, and images were taken using confocal microscopy. For each condition, at least 100 cells were scored. Data are presented as mean ± SEM of three independent experiments. N.D., not determined due to the absence of recombinant protein expression. *, represents significant reduction in nuclear FLAG signal compared with the cells expressing FLAG-NFAT5_132-581_, p<0.0001, by one-way ANOVA with Bonferroni’s multiple comparison test as post-test. Bottom, representative fluorescence images of fixed HeLa cells expressing indicated recombinant NFAT5 mutants under hypotonicity, isotonicity and hypertonicity. Scale bar is 30 µm. **D)** Quantitative analysis of subcellular localization of fluorescence signal in HeLa cells transfected with FLAG-NFAT5_132-581_ and the indicated siRNA from the exportin family. Cells were switched to isotonic or hypotonic medium for 90 mins before fixation. Cells were stained with FLAG antibody and FITC-labeled secondary antibody, counterstained with DAPI. For each condition, at least 100 cells were scored. Data are presented as mean ± SEM of three independent experiments. *, p<0.01, and **, p<0.001 by one-way ANOVA with Bonferroni’s multiple comparison test as post-test, represents significant induction in nuclear fluorescence signal compared with the untransfected cells and cells transfected with non-targeting siRNA. Hypo, hypotonic condition; Iso, isotonic condition. **E)** Representative fluorescence images of endogenous NFAT5, XPOT and XPO4 localization in hypotonic, isotonic and hypertonic medium. HeLa cells were treated with the indicated tonicity for 90 min. NFAT5, XPOT and XPO4 were visualized with corresponding antibodies. Cells were counterstained with Hoechst. Scale bar is 30 µm.

We made deletion mutants (Fig. 4B and Supplementary Fig. 1D) to define the sequence requirement for NFAT5-NLS activity *in vitro*. Compared with His-NFAT5171-250-AcGFP, neither the deletion of amino acid residues 221-250 (His-NFAT5_171-220_-AcGFP) nor amino acid residues 171-189 (His-NFAT5_190-250_-AcGFP) affects KPNB1-mediated nuclear import of the fusion protein (Fig. 4B, lanes 2, 3 and 5). However, deletion of amino acid residues 171-219 (His-NFAT5_220-250_-AcGFP) completely abolished nuclear entry (Fig. 4B, lane 6). Fusion protein containing mainly the cNLS (His-NFAT5_190-230_-AcGFP) failed to undergo nuclear entry (Fig. 4B, lane 4). All fusion proteins failed to enter nucleus in the absence of KPNB1 (Fig. 4B). Collectively, these data corroborated with the result from cell assays that a.a. 189-240 is required for nuclear import activity of NFAT5, and unequivocally demonstrated that Kpnb1 is the only nuclear transport receptor required for the nuclear import.

To identify the critical amino acid residues essential for nuclear import activity *in vivo*, we conducted alanine-scanning mutagenesis (between a.a. 171 to 198, and 228-249) using FLAG-NFAT5_132-581_ as template. We found that alanine substitution of M174, W175, N182, F183, N192, N194, or E196 (proximal to the cNLS), and P233, H234, Y235, L237, or L240 (distal to the cNLS) significantly impaired nuclear localization of NFAT5 under hypertonicity (Fig. 4C), suggesting that these amino acid residues are critical for establishing interaction with KPNB1 for the nuclear import of NFAT5.

### Identification of NTR for NFAT5 nuclear export under hypotonicity

In an earlier study we showed that exportin-1 (CRM1/XPO1) is required for nuclear export of NFAT5 during isotonic nucleocytoplasmic shuttling via acting on the canonical NES(Tong et al., 2006), whereas AED is required for its nuclear export under hypotonicity, although the NTR involved remains unknown(Tong et al., 2006). To identify the putative NTR involved, we conducted siRNA knockdown of exportin genes, followed by determining the subcellular localization of FLAG-NFAT5_132-581_ (which does not have NES, but contains an intact AED) in response to hypotonicity. We successfully knocked down the expression of Exportin-T (XPOT), Exportin-1 (XPO1), Exportin-2 (XPO2), Exportin-4 (XPO4), Exportin-5 (XPO5), Exportin-6 (XPO6) and Exportin-7 (XPO7) (Supplementary Fig. 2A). XPO2 is responsible for nuclear export of KPNAs(Solsbacher et al., 1998), but knockdown of XPO2 has been shown to impact on protein nuclear import (Dong et al., 2018). Consistent with this notion, gene knockdown of XPO2 resulted in exclusive cytoplasmic localization of FLAG-NFAT5_132-581_ under both isotonic and hypotonic conditions (Fig. 4D). Among others, gene knockdown of XPOT or XPO4 mitigated hypotonicity-induced nuclear export of the FLAG-NFAT5_132-581_ (Fig. 4D and Supplementary Fig. 2B). Interestingly, we found that XPOT, but not XPO4, undergoes distinct cytoplasmic and nuclear localization similar to that of NFAT5 in response to hypotonic and hypertonic treatment (Fig. 4E), suggesting that XPOT may be a potential exportin for NFAT5 nuclear export. However, co-immunoprecipitation with FLAG antibodies in cells expressing FLAG-NFAT5_132-581_ failed to identify endogenous XPOT or XPO4 in the immunocomplex (Supplementary Fig. 2C), presumably because of the known transient nature of the exportin-cargo interactions.

### Identification and characterization of novel NFAT5-interacting proteins for nuclear export

The N-terminal region of NFAT5 is associated with many proteins (DuMond et al., 2016). To identify novel NFAT5-interacting proteins that might serve as potential NTRs or regulator(s) for its nuclear export, we treated HeLa cells expressing FLAG-NFAT5_132-264_ with hypotonic and hypertonic medium respectively, followed by affinity-purification using FLAG antibodies and mass spectrometric analysis of the immunocomplexes (Fig. 5A). A total of 163 putative NFAT5 interactors were identified. Among them, 68 and 41 proteins were specifically associated with NFAT5 under hypotonic and hypertonic conditions respectively, whereas 95 proteins were associated with NFAT5 under both conditions (common interactors) (Fig. 5B and Supplementary Table 1).

**Figure 5.**
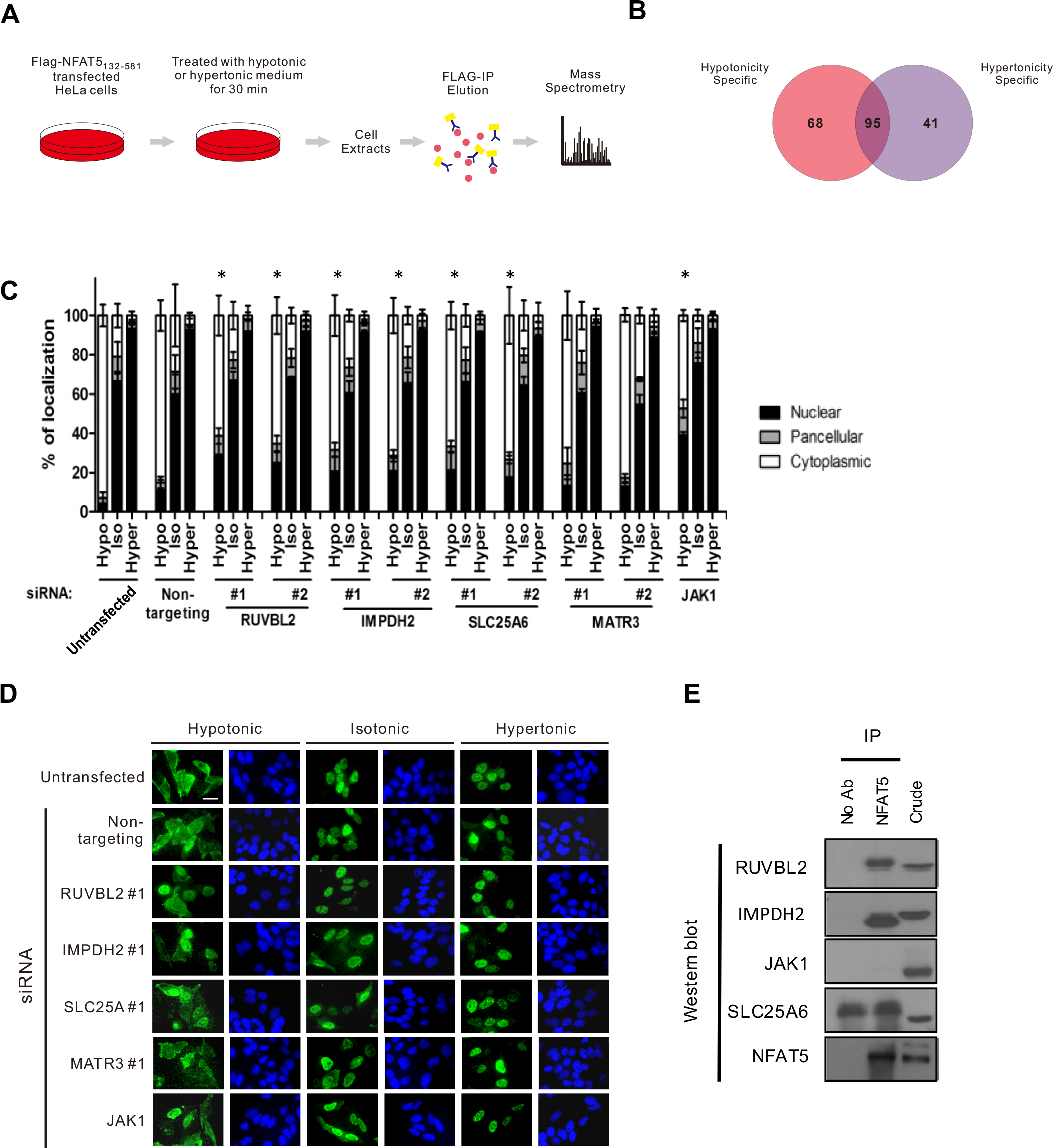
**A)** Schematics of the experimental set up for proteomic identification of NFAT5-interacting proteins. HeLa cells expressing FLAG-NFAT5_132-581_ were switched to hypertonic or hypotonic medium for 30 mins before harvest. Cell extracts were subjected to immunoprecipitation using anti-FLAG antibodies. The immunocomplexes were digested with trypsin and Lys-C, and subjected to mass spectrometric analysis. The MS/MS spectra were subjected to SEQUEST for protein identification. **B)** Venn diagram showing the number of proteins specifically associated with NFAT5 under hypotonic and hypertonic condition respectively, and the number of proteins associated with NFAT5 under both conditions. **C)** Quantitative analysis of subcellular localization of fluorescence signal in HeLa cells transfected with FLAG-NFAT5_132-581_ and the indicated siRNA. Cells were switched to hypotonic, isotonic or hypertonic medium before fixation. For each condition, at least 100 cells were scored. Data are presented as mean ± SEM of three independent experiments. *, represents significant reduction in cytoplasmic fluorescence signal compared with the non-targeting siRNA-expressing cells under the same tonicity, p<0.05, by one-way ANOVA and with Bonferroni’s multiple comparison test as post-test. Hypo, hypotonic condition; Iso, isotonic condition; Hyper, hypertonic condition. **D)** Representative fluorescence images of HeLa cells co-transfected with FLAG-NFAT5_132-581_ and the indicated siRNA. Cells were switched to hypertonic, isotonic or hypotonic medium for 90 mins before fixation. Scale bar is 30 µm. Cells were stained with FLAG antibody and FITC-labeled secondary antibody, counterstained with DAPI. Cells were counterstained with DAPI. **E)** Co-immunoprecipitation analysis of the putative NFAT5-interacting proteins. Immunoprecipitation was carried out using NFAT5 antibodies and the immunocomplexes were subjected to Western blot analysis using the indicated antibodies.

A total of ninety-five putative NFAT5 interactors (hypotonicity-specific and common interactors) were selected for functional evaluation for their role in hypotonicity-induced NFAT5 nuclear export. Cytoskeletal proteins, histones, ribosomal proteins, and mitochondrial proteins were excluded in the analysis because it is known that these proteins tend to associate with the immunocomplex in a non-specific manner. SMARTpool siRNAs were used to knock down the expression of each of these proteins, and subcellular localization of FLAG-NFAT5_132-264_ under different tonicities was determined respectively. We found that cells expressing siRNAs against RuvB-Like AAA type ATPase (REPTIN or RUVBL2), Inosine-5’-monophosphate dehydrogenase 2 (IMPDH2), Janus kinase 1 (JAK1), Adenine Nucleotide Translocator 3 (SLC25A6), and Matrin 3 (MATR3) profoundly inhibited cytoplasmic localization of FLAG-NFAT5_132-264_ under hypotonic condition (defined as % of localization_cytoplasmic_ < 50) (Supplementary Fig. 3).

Secondary screening of each putative interactor using two independent siRNAs (Supplementary Fig. 4A) confirmed that hypotonicity-induced nuclear export of FLAG-NFAT5_132-264_ was significantly blocked by gene knockdown of RUVBL2, SLC25A6, IMPDH2, or JAK1 (one siRNA) (Fig. 5C and 5D). Furthermore, immunoprecipitations showed that endogenous NFAT5 is associated with RUVBL2 and Impdh2 respectively, but not with SLC25A6 or JAK1 (Fig. 5E). The role of RUVBL2 was further characterized in this study, because we observed a distinctive nuclear accumulation of the FLAG-NFAT5_132-264_ when RUVBL2 was depleted (Fig. 5D).

### RUVBL2 is required for hypotonicity-induced NFAT5 nuclear export

RUVBL2 and RUVBL1, which are also known as reptin and pontin respectively, are two closely related proteins belonging to the large AAA+ ATPase (ATPase associated with diverse cellular activities) superfamily that are characterized by an ATPase core domain with Walker A and Walker B motifs (Jha and Dutta, 2009). RUVBL2 and RUVBL1 can either complex with each other and function together (Izumi et al., 2010), or act independently from each other (Bauer et al., 2000; Rottbauer et al., 2002; Diop et al., 2008; Kim et al., 2005) to regulate a wide variety of functions including chromatin remodeling, transcription regulation, DNA damage response, and assembly of ribonucleoprotein complexes(Izumi et al., 2010). We found that the expression of siRNA-resistant RUVBL2 in cells transfected with RUVBL2 siRNA restored hypotonicity-induced nuclear export of FLAG-NFAT5_132-581_ (Fig. 6A). RUVBL2 contains a ATPase domain that may be essential for its function(Grigoletto et al., 2011). Expression of siRNA-resistant RUVBL2 ATPase mutant, RUVBL2 (E300G), restored NFAT5 nuclear export similar to the overexpression of siRNA-resistant RUVBL2 (Fig. 6A and Supplementary Fig. 4B). Concordantly, we found that CB-6644, a small-molecule inhibitor of the ATPase activity of the RUVBL1/2(Assimon et al., 2019), failed to block hypotonicity-induced nuclear export of FLAG-NFAT5_132-264_ (Supplementary Fig. 4C). Our findings suggested that RUVBL2 regulate NFAT5 nuclear export in an ATPase-independent manner. On the other hand, siRNA knockdown of RUVBL1 did not significantly alter NFAT5 nuclear export (Supplementary Fig. 4D), suggesting that RUVBL2 did not cooperate with RUVBL1 to mediate the process. Furthermore, co-immunoprecipitation suggested that FLAG-NFAT5_132-264_ and RUVBL2 was consistently associated with each other, but the interaction between the two proteins was reduced under hypertonicity (Fig. 6B). Similarly, we observed reduced association between endogenous NFAT5 and RUVBL2 under hypertonic condition (Fig. 6C), which was not due to reduced availability of RUVBL2 under hypertonicity (Supplementary Fig. 4E).

**Figure 6.**
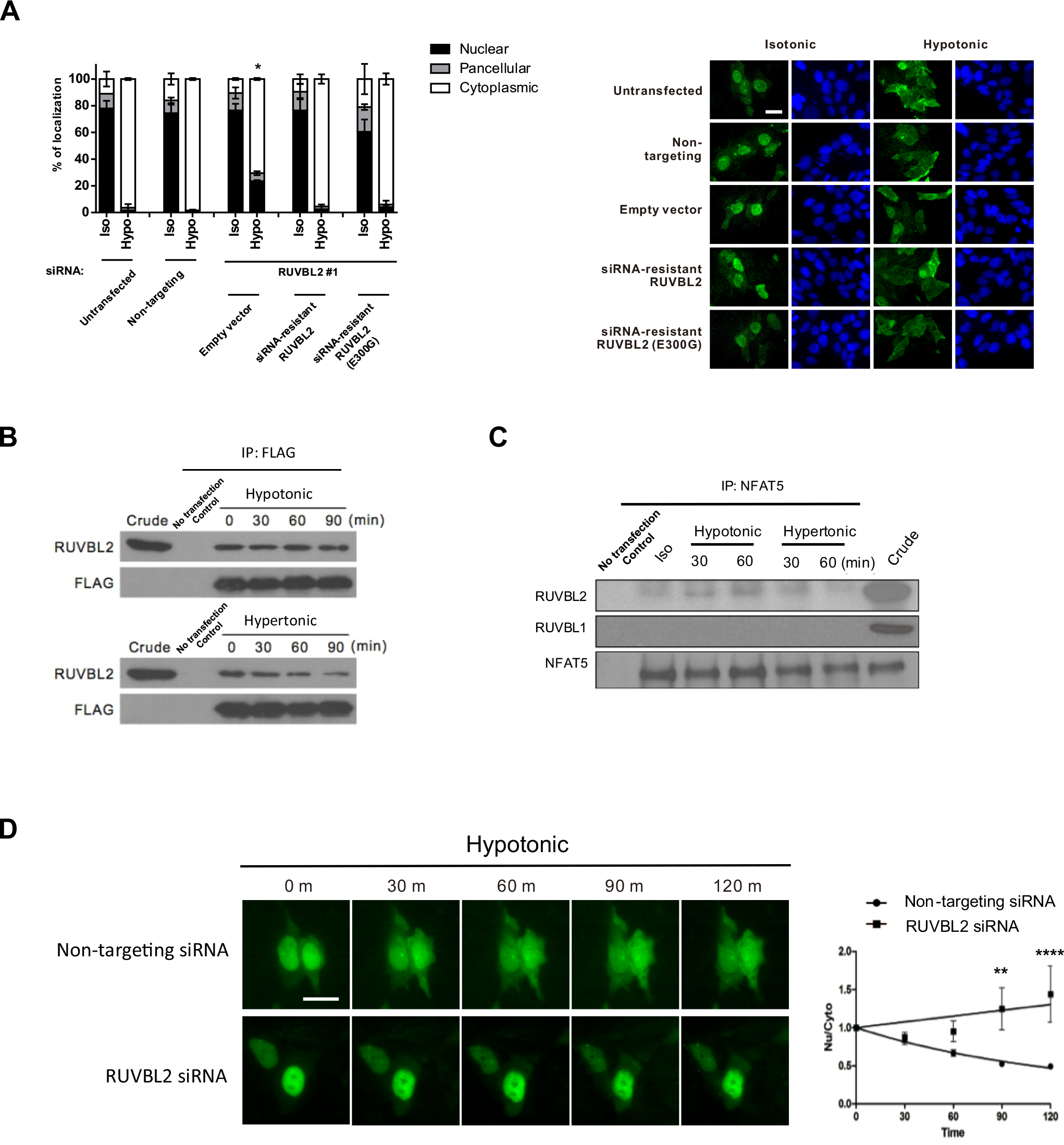
**A)** Rescue of RUVBL2 expression restored NFAT5 nuclear export. Quantitative analysis of subcellular localization of FLAG-NFAT5_132-581_ in cells co-transfected with RUVBL2 siRNA and siRNA-resistant wild type RUVBL2 (WT) or RUVBL2 E300G mutant. Cells were switched to isotonic or hypotonic medium for 90 mins before fixation. For each condition, at least 100 cells were scored. Data are presented in mean ± SEM of two independent experiments. *, represents significant induction in nuclear fluorescence compared with that of the non-targeting siRNA-expressing cells (hypo), p<0.01 by one-way ANOVA with Bonferroni’s multiple comparison test as post-test. Iso, isotonic condition; Hypo, hypotonic condition. Representative fluorescence images were shown, scale bar is 30 µm. **B)** Interaction between NFAT5 and RUVBL2. Cell extracts prepared from HeLa cells expressing FLAG-NFAT5_132-581_ treated with the indicated tonicity were immunoprecipitated with FLAG antibodies. The immunocomplexes were subjected to Western blot using the RUVBL2 antibodies. **C)** NFAT5 was associated with RUVBL2 under isotonic and hypotonic condition. HeLa cells extracts prepared from cells treated with the indicated tonicity were immunoprecipitated with NFAT5 antibodies. The immunocomplexes were subjected to Western blot hybridization using the RUVBL2 and RUVBL1 antibodies. **D)** Live cell monitoring of NFAT5-EGFP fusion protein trafficking in response to hypotonic treatment. HeLa cells were transfected with NFAT5_128-581_-EGFP and non-targeting siRNA (Control), or NFAT5_128-581_-EGFP and RUVBL2 siRNA (RUVBL2) respectively. Cells were pre-treated with cycloheximide for 1 h and then induced with hypotonic medium and EGFP fluorescence was captured at 0, 30, 60, 90 and 120 min. Left, representative images of GFP fluorescence, scale bar is 30 µm. Right, quantification of nuclear (Nu) / Cytoplasmic (Cyto) fluorescence signal in cells transfected with non-targeting siRNA vs RUVBL2 siRNA. Data are presented in mean ± SEM of three independent experiments (For each condition, n≥9 cells were analyzed). **, p < 0.01, ****, p < 0.0001, by two-way ANOVA with Bonferroni’s multiple comparison test as post-test.

To further substantiate the function of RUVLB2 in NFAT5 nuclear export, we expressed a NFAT5-EGFP reporter protein (NFAT5_128-581_-EGFP) for real time monitoring of its trafficking in cells expressing non-targeting siRNA and RUVBL2 siRNA respectively, using time-lapse fluorescence photography as we have described(Xu et al., 2008). We found that hypotonicity induced time-dependent nuclear export of GFP signal in cells expressing non-targeting siRNA, but the export was remarkably mitigated in cells expressing RUVBL2 siRNA (Fig. 6D). Together, these data suggested that RUVBL2 is essential for nuclear export under hypotonicity.

### Nucleocytoplasmic trafficking of RUVBL2 in response to changes in extracellular tonicity

RUVLB2 primarily resides in the nucleus(Bauer et al., 1998; Sigala et al., 2005). We first determined its subcellular localization under different extracellular tonicities. Endogenous RUVBL2 is localized pan-cellularly under isotonic condition. It becomes predominantly localized to the cytoplasm and nucleus in response to hypotonicity and hypertonicity challenge respectively (Fig. 7A), suggesting that RUVBL2 undergoes nucleocytoplasmic trafficking in response to changes in extracellular tonicity *per se*. To further understand the relationship between RUVBL2 and NFAT5, we created a Dendra2-RUVBL2 reporter construct (Fig. 7B). Dendra2 is a green-emitting fluorescent protein that can be converted to emit red light upon excitation at 405-nm (Gurskaya et al., 2006). Upon photoconversion, it allows live-cell monitoring of the converted protein without interference from newly synthesized or unconverted proteins (Fig. 7B)(Chudakov et al., 2007). Similar to the endogenous RUVBL2, Dendra2-RUVBL2 is localized pan-cellularly (Fig. 7C, green fluorescence). Time-lapsed confocal photography showed that, in cells transfected to non-targeting siRNA, photoconverted Dendra2-RUVBL2 remains residing in the nucleus at 60 min under isotonicity. Under hypertonic condition, it undergoes cytoplasmic translocation in a time-dependent manner, with a concomitant reduction of signal intensity (Figs. 8C and 8D). Remarkably, hypotoncity-induced nuclear export of photoconverted Dendra2-RUVBL2 was not affected by reduced availability of NFAT5 by transfection of NFAT5 siRNA (Figs. 8C, 8E, and 8F). Taken together, our data suggested that RUVBL2 is required for hypotonicity-induced nuclear export of NFAT5, but not vice versa. Therefore, RUVBL2 is a chaperone for NFAT5 nuclear export.

**Figure 7.**
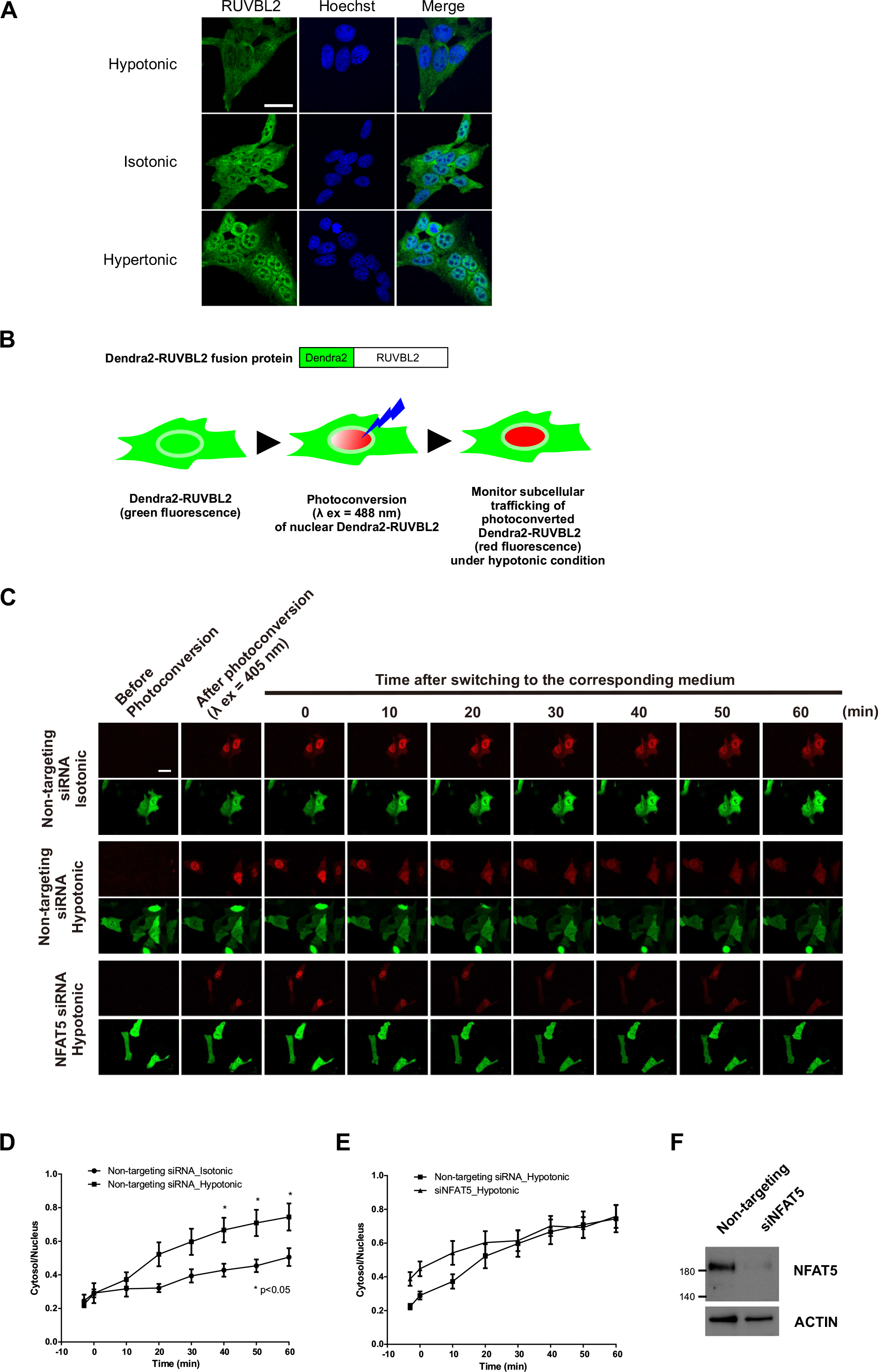
Nucleocytoplasmic trafficking of RUVBL2 and NFAT5. **A)** Representative fluorescence images of endogenous RUVBL2 localization in hypotonic, isotonic and hypertonic medium. HeLa cells were treated with the indicated tonicity for 90 min. RUVBL2 was visualized with a RUVBL2 antibody. Cells were counterstained with DAPI. Scale bar is 30 µm. **B)** Schematics of the Dendra2-RUVBL2 experimental setup. HeLa cells expressing Dendra2-RUVBL2 and non-targeting siRNA or NFAT5 siRNA respectively were photoconverted with 405nm laser. Dendra2-RUVBL2 in the nucleus was photoconverted from green to red. The Dendra2-RUVBL2 protein trafficking in live cells in response to hypotonic treatment was monitored. **C)** Dendra2-RUVBL2 time-lapse experiment of the HeLa cells transfected with non-targeting siRNA or NFAT5 siRNA. Top, representative red fluorescence images of photoconverted Dendra2-RUVBL2 fusion protein. Bottom, representative green fluorescence images of non-photoconverted Dendra2-RUVBL2 fusion protein in cells. After 405 nm photoconversion, cells were treated with hypotonic medium. Fluorescence images were captured every 10 min. Scale bar is 30 µm. **D)** the quantification of cytoplasmic (Cyto) / nuclear (Nu) red fluorescence signals in cells transfected with non-targeting siRNA or **E)** NFAT5 siRNA respectively under hypotonic stress. Data are presented in mean ± SEM of three independent experiments (For each condition, n≥10 cells were analyzed). *, p < 0.05, by two-way ANOVA with Bonferroni’s multiple comparison test as post-test. **F)** Western blot analysis of NFAT5 expression in cells transfected with non-targeting and siNFAT5 siRNAs.

**Figure 8.**
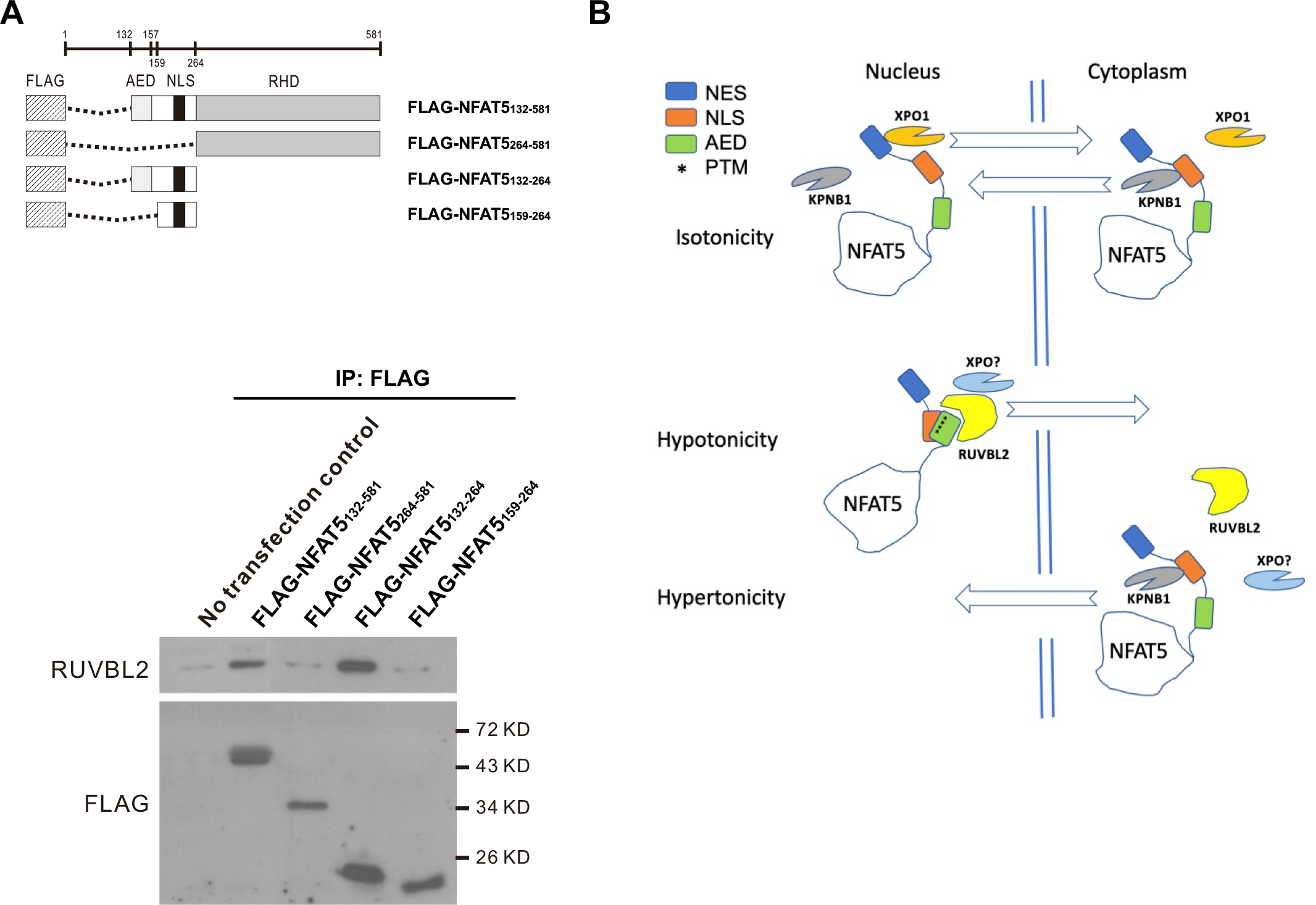
**A)** Top, schematics of FLAG-NFAT5 deletion constructs. Bottom, HeLa cells expressing the indicated NFAT5 deletion mutant were immunoprecipitated using anti-FLAG affinity resin. The immunocomplex was subjected to Western blot analysis using RUVBL2 antibodies. **B)** Schematics showing the proposed mechanisms of the nuclear import and nuclear export of the NFAT5 under different extracellular tonicities.

### Interactions between NFAT5 and RUVBL2

To further understand if and how NFAT5 and RUVBL2 interact with each another, we expressed FLAG-NFAT5 deletion mutants (Fig. 8A, top) for co-immunoprecipitation analysis. As expected, co-immunoprecipitation of FLAG-NFAT5_132-581_ resulted in the presence of endogenous RUVBL2 in the immunocomplex. However, RUVBL2 signal was abolished in the immunocomplex of FLAG-NFAT5_264-581_ (Fig. 8A, bottom). Similarly, RUVBL2 was associated with FLAG-NFAT5_132-264_, but it failed to form any complexes with FLAG-NFAT5_159-264_ (Fig. 8A, bottom). These data suggested that AED is crucial for NFAT5 to interact with RUVBL2, and hence NFAT5 nuclear export. To further determine if RUVBL2 interacts with AED directly, we expressed recombinant FLAG-NFAT5131-249 and His-RUVBL2 and tested their binding by ITC assay. However, we failed to detect any interaction even after repeated trials (Supplementary Fig. 4F), suggesting that the two proteins may not interact directly under this setting.

## Discussion

NFAT5 was discovered two decades ago as the first transcription factor involved in the orchestrating cellular adaptive responses to hypertonic stress(Ko et al., 2000; Miyakawa et al., 1999), and it is the best characterized tonicity-regulated transcription factor to date. Hypertonic stress was once considered to be present only in the inner medulla of the kidneys (Burg et al., 2007), but emerging evidence has suggested that excessive sodium accumulation in other tissue microenvironments, such as the skin (Jantsch et al., 2015) and the intervertebral disc (Switzerland et al., 2018), may activate NFAT5 that leads to either protective (Machnik et al., 2009; Switzerland et al., 2018) or pathogenic (Matthias et al., 2019; Ma et al., 2020) outcomes. Although the contemporary view on how sodium accumulation leads to NFAT5 activation in these tissues has been challenged recently (Rossitto et al., 2020), there is evidence suggesting that dysregulation of NFAT5 contributes to the pathogenesis of autoimmune diseases(Matthias et al., 2019; Aramburu and Lopez-Rodriguez, 2019; Lee et al., 2019; Kleinewietfeld et al., 2013), inflammation(Aramburu and Lopez-Rodriguez, 2019), hypertension and metabolic disorders(Jantsch et al., 2014).

While the physiological and pathological role of NFAT5 has been firmly established, the molecular mechanisms underlying how NFAT5 activity is regulated remain largely elusive. Our study has generated multiple insightful findings into tonicity-regulated trafficking mechanisms of NFAT5 with both mechanistic and therapeutic implications. First, we provided direct evidence of visualizing NFAT5 trafficking through the nuclear pore complex by using NFAT5-MiniSOG fusion protein assay and visualized it by electron microscopy. This novel assay allowed the observation of the “double ring” conformation of NFAT5 under hypertonicity *in vivo*, which convincingly supported a previously proposed homodimeric structural model for NFAT5 (Stroud et al., 2002). Together these findings strengthen the notion that NFAT5 homodimer binds to target DNA through complete encirclement of the DNA(Stroud et al., 2002). Interestingly, the ring-like conformation of NFAT5 was not observed under hypotonicity, suggesting that NFAT5 exits as monomer in the cytoplasm. Our nucleocytoplasmic trafficking analysis using reporter constructs which lack RHD and DD further confirms that NFAT5 could undergo nucleocytoplasmic trafficking in monomeric form.

A previous bioinformatics analysis has suggested that NFAT5 contains putative protein domain exhibiting high sequence homology to monopartite cNLS (Tong et al., 2006), and it is presumed that its nuclear import is mediated by a heterotrimeric Kapα/Kapβ/NFAT5 complex (Lott and Cingolani, 2011). Unexpectedly, we found that NFAT5 utilizes an unconventional mechanism for nuclear import in which KPNB1 alone serves as both the adaptor and the importin. KPNB1-driven NFAT5 nuclear import requires the interaction between KPNB1 and an extended region of NFAT5-NLS. How the NFAT5-NLS region interacts with KPNB1 remains to be determined. Previous studies have shown that the ncNLSs can adopt a variety of secondary structures such as α helices in importin-α-IBB (Cingolani et al., 1999) and SREBP2 (Lee et al., 2003), the C2H2 zinc finger domain in Snail (Choi et al., 2014) and the long and unstructured NLS of hnRNP A1 (Lee et al., 2006a). On the other hand, KPNB1 shows conformational flexibility across its elongated superhelical structure, allowing it to effectively “wrap” around structurally diverse ncNLSs for tight and specific interaction. Accordingly, the binding sites on KPNB1 for these ncNLSs vary from the N-terminal ten HEAT repeats for parathyroid hormone-related protein to the last few C-terminal HEAT repeats for SREBP2 (Cingolani et al., 2002). For NFAT5, although the short NFAT5-cNLS sufficiently established interaction with KPNB1, nuclear import activity could only be elicited by the extended NFAT5-NLS. Our results suggested that the long and unstructured NFAT5-NLS segment may fit onto the concave surface of KPNB1 for specific and direct interaction. Such interaction may be required for inducing conformational changes of KPNB1 to elicit its nuclear import activity. Structural elucidation of the NFAT5-NLS-KPNB1 complex by crystallographic analysis in the future may provide a better understanding regarding the underlying mechanism. Members of the NFAT family (NFAT1-5) are characterized by the presence of Rel homology DNA-binding domains (Graef et al., 2001). It is hypothesized that distinctive genome recombination events during evolution had led to differential acquisition of different regulatory sequences, giving rise to tonicity-responsive NFAT5, and calcium-responsive NFAT1-4(Graef et al., 2001) that are involved in diverse cellular functions ranging from T cell response to organogenesis(Zhu and McKeon, 2000). NFAT5 homolog can be found in drosophila, whereas NFAT1-4 is restricted to vertebrates, suggesting that NFAT5 has a more ancestral origin(Graef et al., 2001). Our finding therefore suggested that the classical nuclear import pathway evolved for nuclear import of NFAT1-4 serves to differentiate activation of NFAT1-4 and NFAT5 activation.

On the other hand, we have identified XPOT and XPO4 as potential exportins for NFAT5 nuclear export. XPOT and XPO4 are known for their role in nuclear export of tRNA(Arts et al., 1998) and translation initiation factor eIF-5A(Lipowsky et al., 2000) respectively. XPOT is particularly interesting because it undergoes nucleocytoplasmic trafficking in response to changes in extracellular tonicity similar to NFAT5, aligning with its role as a NFAT5 exporter. Our findings further suggested that nuclear export of NFAT5 might be coupled with tRNA export. More importantly, we have identified RUVBL2 as a novel regulator for NFAT5 nuclear export under hypotonicity. In corroboration with our findings, a recent proteomic analysis of NFAT5-interacting proteins has also identified RUVBL2 in the NFAT5 immunocomplex(DuMond et al., 2016). RUVBL2 (also known as Reptin) and RUVBL1 (also known as Pontin) are two closely related members of AAA+ family of DNA helicases closely related to bacterial DNA helicase RuvB, a member of the AAA+ ATPase family of helicases (<u>A</u>TPase <u>a</u>ssociated with diverse cellular <u>a</u>ctivities) (Ogura and Wilkinson, 2001). RUVBL1 and RUVBL2 were found to exist as monomer, homodimer, and homo- or hetero-hexamer (Niewiarowski et al., 2010). Although the role and regulation of ATP hydrolysis remain unclear, RUVBL1-RUVBL2 hetero-hexameric complex acts as a scaffold for protein-protein interactions (Dauden et al., 2021). We found that RUVBL2 mediates NFAT5 nuclear export in a novel manner that does not require ATPase activity or RUVBL1. Interestingly, we found RUVBL2 *per se* is subjected to tonicity-dependent nucleocytoplasmic trafficking in response to changes in tonicity. NFAT5 nuclear export was inhibited when RUVBL2 was depleted, but not vice versa, suggesting that RUVBL2 may act as a hypotonicity-specific chaperone of NFAT5 for its nuclear export. Taken together, similar to the role RUVBL1-RUVBL2 complex as a protein scaffold(Dauden et al., 2021), RUVBL2 might bridge NFAT5 to XPOT for nuclear export.

Nuclear availability of many transcription factors is regulated by differential exposure of NLS and NES respectively. For example, in response to calcium signaling, NFAT1 nuclear import is regulated by dephosphorylation of the regulatory domain, resulting in the masking of NES and exposure of NLS(Okamura et al., 2000). We proposed that NFAT5 activity may be regulated in a similar manner. We proposed a mechanistic model in Fig. 10B. Under isotonicity, NFAT5-NLS collaborates with the classical NES (a.a. 8-15) through binding to KPNB1 and XPO1 respectively, for maintaining NFAT5 nucleocytoplasmic shuttling in a homeostatic state. AED and NLS are located in close proximity within the intrinsically disordered (ID) region, which is well-known for its structural flexibility for differential interaction with proteins(Garza et al., 2009). A number of tonicity-dependent PTMs critical for nucleocytoplasmic trafficking of NFAT5, such as phosphorylation of T135 and Y143 (Gallazzini et al., 2011, 2010), as well as S155 and S188 (Tong et al., 2009), are located within this region. Specific PTMs induced by hypotonicity might induce conformational changes in NFAT5, resulting in the masking of NLS and the association with RUVBL2 and XPOT for nuclear export. In contrast, specific PTMs associated with hypertonicity might lead to NLS exposure for nuclear import. Accordingly, the lack of appropriate PTMs in the recombinant NFAT5 proteins or the absence of XPOT might account for the failure to detect any direct interaction between NFAT5 and RUVBL2 in the *in vitro* ITC analysis. On the other hand, under hypertonicity, reverse conformational changes might lead to preferential binding of NFAT5 to KPNBP1, resulting in net nuclear import (Fig. 8B).

In conclusion, we have discovered and characterized a number of novel players responsible for tonicity-regulated nucleocytoplasmic trafficking of NFAT5. Tonicity-dependent NFAT5 activation has been associated with a variety of physiological processes and diseases, including blood pressure regulation, inflammation, and development of autoimmune diseases (Machnik et al., 2009; Kleinewietfeld et al., 2013). Understanding the mechanisms of its nucleocytoplasmic trafficking mechanisms may lead to the development of therapeutic strategies for diseases that are associated with NFAT5 dysregulation.

## Notes

**Funding** This work was supported by RGC General Research Fund (15106417), PolyU internal grant (P0009343), Research Impact Funds (R5050-18 and R4015-19F), and Collaborative Research Fund Equipment Grant (C5012-15E), NIH R01GM086197 (DB), and NIH P41 GM103412 for support of the NCMIR (MHE).

### Competing Interest Statement

The authors have declared no competing interest.

