## Supplementary Fig1-4 for "Unconventional tonicity-regulated nuclear trafficking of NFAT5 mediated by KPNB1, XPOT and RUVBL2"

**Supplementary Legends**

**Supplementary Figure 1. A)** Western blot analysis of FLAG-NFAT5$_{132-581}$ and FLAG-NFAT5-PEPCK mutants. **B)** Representative fluorescence images of FLAG-NFAT5$_{132-581}$ in the cells expressing the KPNB1 siRNAs. Cells were counterstained with DAPI. Scale bar is 30 μm. **C)** GST-NFAT5 proteins were purified using glutathione sepharose column. **D)** Recombinant His-NFAT5-AcGFP proteins were purified using HisTrap column. Purified proteins in C and D were subjected to SDS-PAGE analysis followed by Coomassie blue staining. All purified proteins were expressed at the expected molecular weight. **E)** *In vitro* nuclear import assay. Nuclear import of His-SV40 NLSAcGFP, His-NFAT5171-250-AcGFP, and His-AcGFP were determined using digitonin permeabilized HeLa cells. Transport assay was carried out for 30 min at 37$^{o}$C, cells were fixed with paraformaldehyde, stained with DAPI and images were taken using confocal microscopy. Scale bar is 30 μm.

**Supplementary Figure 2. A)** Quantitative real-time PCR analysis of the exportins siRNA knockdown efficiency. Data are presented in mean ± SEM; *, $p < 0.05$, by unpaired t-test. **B)** Representative fluorescence images of HeLa cells transfected with FLAG-NFAT5$_{132-581}$ and the indicated siRNA from the exportin family. Scale bar represents 30 μm. Cells were switched to isotonic or hypotonic medium for 90 mins before fixation. Cells were stained with FLAG antibody and FITC-labeled secondary antibody, counterstained with DAPI. **C)** Interaction between NFAT5 and XPOT/4. HeLa cells expressing FLAG-NFAT5$_{132-581}$ prepared from cells treated with hypotonicity for 30 min were immunoprecipitated with FLAG antibodies. The immunocomplexes were subjected to Western blot hybridization using the XPOT and XPO4 antibodies.

**Supplementary Figure 3.** Quantification of the subcellular localization of FLAG-NFAT5$_{132-581}$ in HeLa cells in response to the expression of the indicated SMARTpool siRNAs. Cells were treated with hypotonic (Hypo), isotonic (Iso) or hypertonic (Hyper) medium for 90 mins. Recombinant protein was visualized with FLAG antibodies and FITC-labeled secondary antibodies, counterstained with DAPI, and analyzed by fluorescence microscopy. For each condition, 100 cells were counted. *, represents significant differences in FLAG signal compared with the cells transfected with non-targeting SMARTpool siRNA.

**Supplementary Figure 4. A)** Western blot analysis and quantitative real-time PCR analysis of the siRNA knockdown efficiency of the indicated putative target genes. Data of qPCR are presented in mean ± SEM; *, $p < 0.05$, **, $p < 0.01$, by one-way ANOVA with Bonferroni's multiple comparison test as post-test. B**)** Western blot analysis indicates rescue of RUVBL2 in cells expressing RUVBL2 siRNA with either siRNA-resistant wild-type (WT) RUVBL2 or siRNA-resistant RUVBL2 ATPase mutant (E300G), respectively. **C)** Quantitative analysis of the subcellular localization of FLAG-NFAT5$_{132\text{-}581}$ in CB-6644 treated cells. Cells were pre-treated with CB-6644 for 2h and then treated with either isotonic or hypotonic medium with CB-6644 respectively. Iso, isotonic condition; Hypo, hypotonic condition. Representative fluorescence images were shown, scale bar is 60 µm. **D)** siRNA analysis of RUVBL1. Top, representative fluorescence images of FLAG-NFAT5$_{132\text{-}581}$ in RUVBL1 siRNA expressing cells. Cells were counterstained with DAPI. Scale bar is 30 µm. Bottom left, Quantitative analysis of siRNA knockdown of RUVBL1 in the subcellular localization of FLAG-NFAT5$_{132\text{-}581}$. Cells were treated with hypotonic, isotonic or hypertonic medium respectively. For each condition, >100 cells were counted. Hypo, hypotonic condition; Iso, isotonic condition; Hyper, hypertonic condition. Bottom right, the quantitative real-time PCR analysis of the siRNA knockdown efficiency of the indicated putative target genes. **E)** Western blot analysis indicates the endogenous expression level of RUVBL2 under different tonic conditions. Cells were treated with hypotonic, isotonic or hypertonic medium for 90 mins. **F)** Isothermal Titration Calorimetry (ITC) profiles to measure the binding affinity of NFAT5 to RUVBL2.

**Supplementary Table 1.** NFAT5 interacting proteins identified by mass spectrometric analysis. Protein candidates in the shaded boxes are either mitochondria, ribosomal, cytoskeletal proteins or histones, and were not included in the siRNA screen.

Supplementary Fig. 1

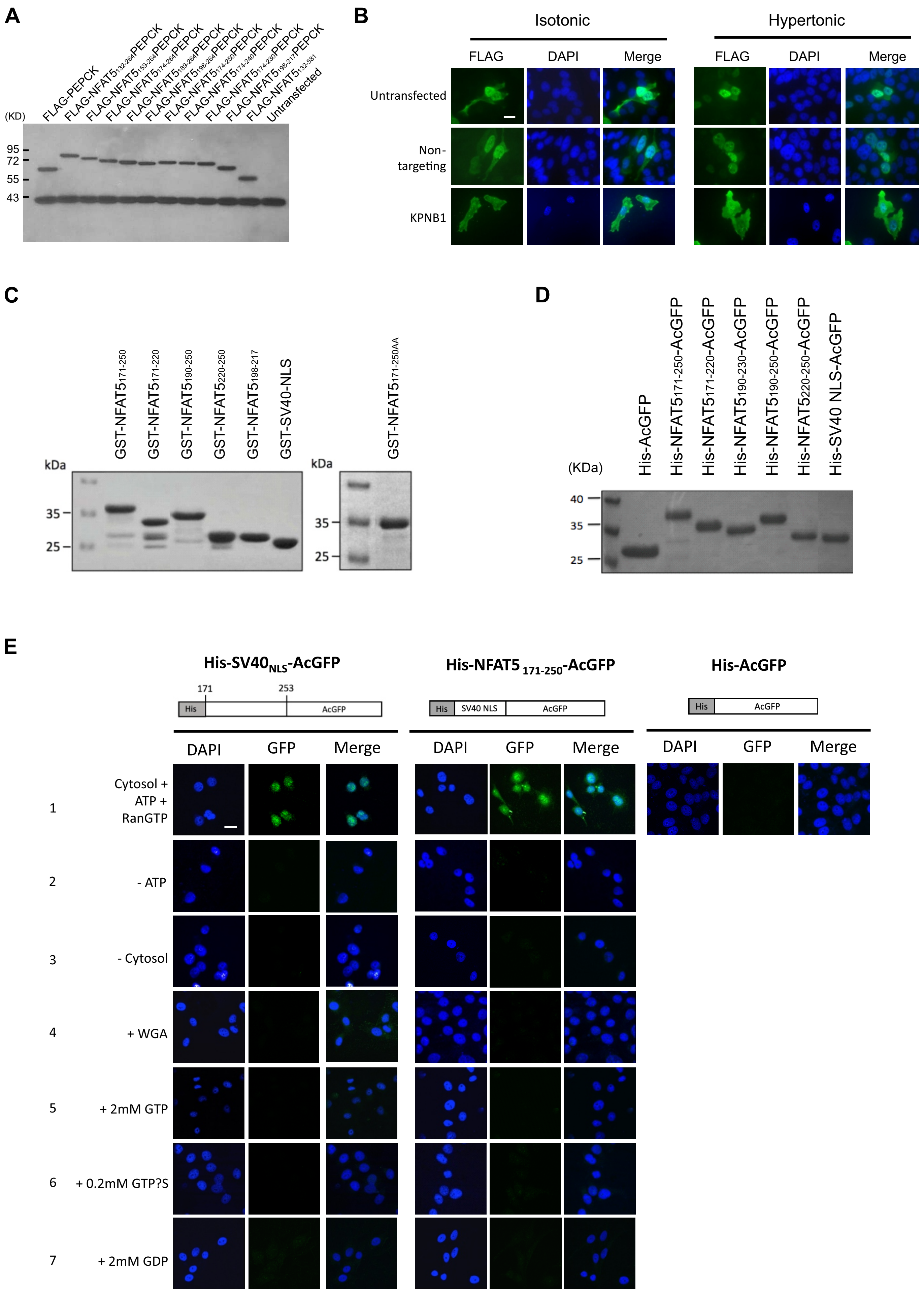

### Supplementary Fig. 2

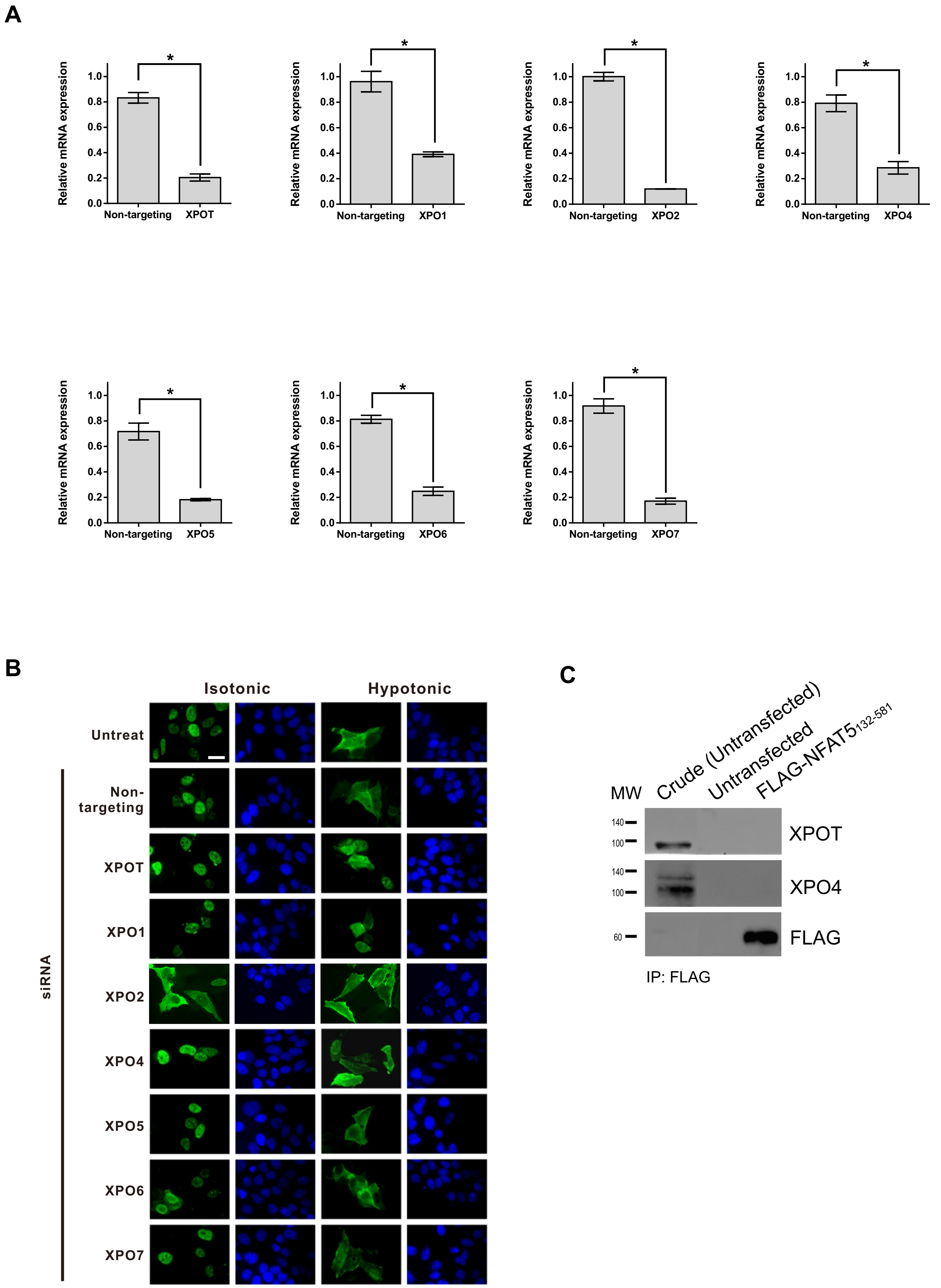

Supplementary Fig. 3

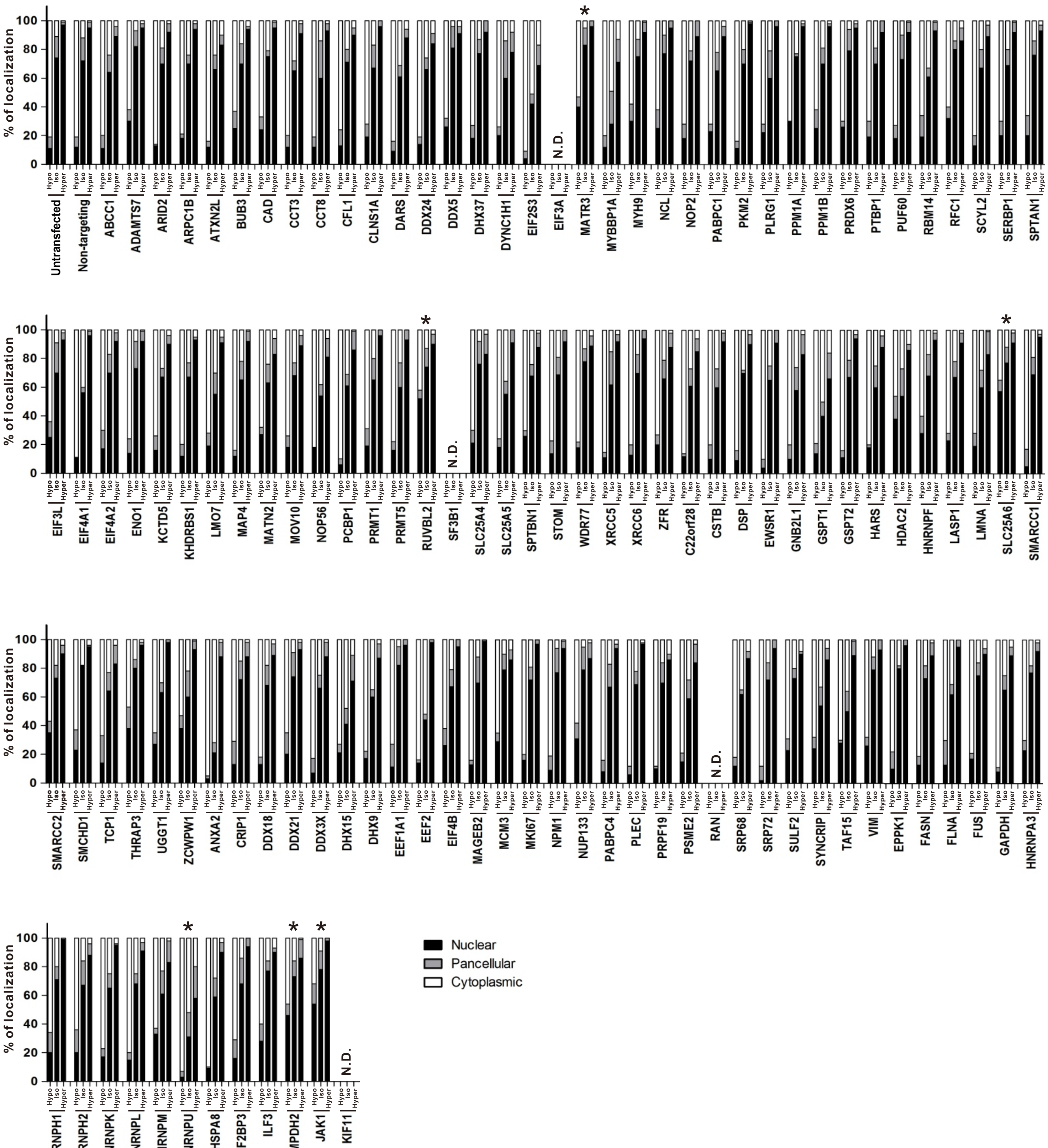

### Supplementary Fig. 4

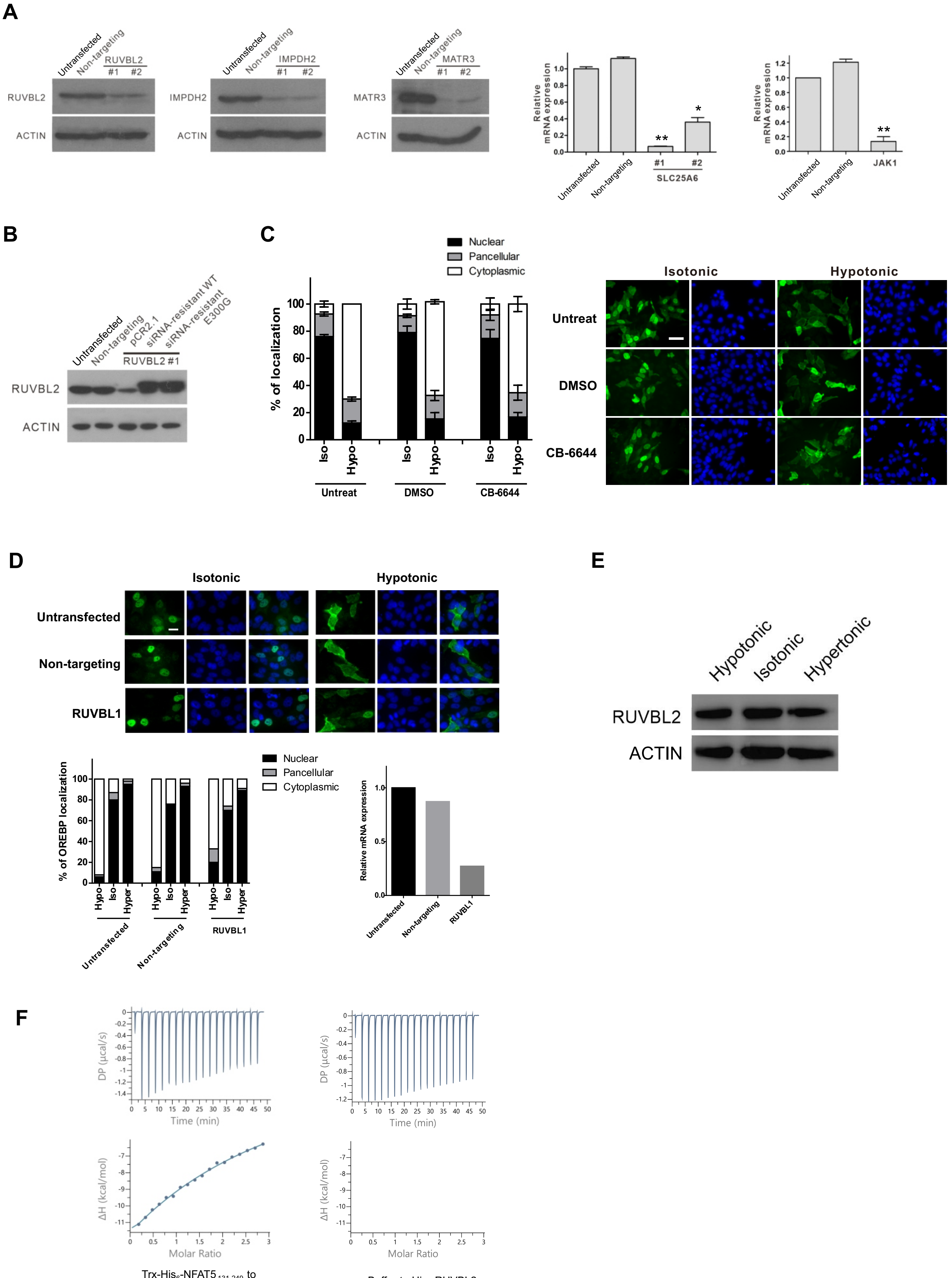


Trx-His$_6$-NFAT5$_{131\text{-}249}$ to His$_6$-RUVBL2

Buffer to His$_6$-RUVBL2
